# PI3Kδ Bridges Microbial Surveillance with Antigen Presentation to Reinforce Intestinal Immunity

**DOI:** 10.1101/2025.05.13.653657

**Authors:** Maria Gonzalez Nuñez, Luiz Ricardo da Costa Vasconcellos, Jose Garrido Mesa, Rafael Cardoso Maciel Costa Silva, Abderrahman Hachani, Alba Bosquet Agudo, Raffaele Simeoli, Laura Medrano Gonzalez, Alain Filloux, Heidi C. E. Welch, Mauro Perretti, Bart Vanhaesebroeck, David Dombrowicz, Oliver Haworth, Klaartje Kok, Bénédicte Manoury, Ezra Aksoy

## Abstract

Phosphoinositide 3-kinase delta (PI3Kδ) is essential for immune cell functions, preventing immunodeficiency and inflammation; however, its role in dendritic cell (DC)-mediated immune regulation remains unknown. Here, we report a key role for DC-intrinsic PI3Kδ in linking microbial recognition with antigen presentation for effective T-cell priming. Using genetic and functional assays, we demonstrate that PI3Kδ deficiency in DCs leads to broad dysregulation of intestinal CD4⁺ T-cell immunity, characterized by impaired regulatory T-cell expansion and increased susceptibility to colitis. DC-based studies show that PI3Kδ links pattern-recognition-receptor signaling to MHC class I- and II-restricted presentation of phagosome-associated antigens by facilitating NOX2-dependent oxidative burst through RAC2, while mitigating inflammasome activation. In contrast, PI3Kδ deficiency disrupts phagosomal pH balance, leading to accelerated acidification and proteolysis of antigens, impairing T-cell activation. Our study identifies PI3Kδ as a key coordinator of phagosome dynamics, important for DC adaptive programming that shapes T-cell responses and supports intestinal immune homeostasis.

## Introduction

Phosphoinositide 3-kinases (PI3Ks) are lipid-modifying enzymes involved in mammalian immune function^1^. The PI3Kδ isoform belongs to the class I PI3K family, comprising three isoforms that include the catalytic subunits p110α, β, γ and δ, complexed with regulatory subunits p85 or p101/p84 as functional heterodimers (also referred as PI3Kα, β, and δ)^2^. The enzymatic activity of the p110 catalytic subunits converts the phosphoinositide (4,5) bisphosphate (PtdIns(4,5)P_2_) lipid to phosphoinositide(3,4,5)triphosphate (PtdIns(3,4,5)P_3_,) a second messenger critical for recruiting pleckstrin homology (PH) domain-containing proteins, notably AKT/PKB to the plasma membrane^3, 4^. p110 subunits also interact with Ras, Rho GTPases and GPCR subunits, while the p85 subunit mediates receptor anchoring via Src homology 2 (SH2) domains that recognize phosphorylated tyrosine (pY) motifs on receptor tyrosine kinases (RTKs) and adaptor proteins^2, 5^.

In leukocytes, PI3Kδ and PI3Kγ are the predominant class I PI3K isoforms, and couple immune receptor signaling to key innate and adaptive immune responses^1, 6^. Inborn errors affecting PI3Kδ expression or function manifest as primary immunodeficiency syndromes, characterized by general immunological dysregulation and heightened susceptibility to opportunistic microbial infections, particularly in the lung and gastrointestinal tract^7, 8, 9^. These genetic insights are paralleled by clinical observations in oncological interventions, where targeted PI3Kδ inhibition frequently precipitates gastrointestinal toxicity, immunosuppressive and inflammatory complications^10, 11, 12^.

Mice carrying an inactivating mutation in the *Pik3cd* gene encoding p110δ catalytic isoform (referred to hereafter as δD910A)^13^, provide a model for exploring intestinal immune regulation. δD910A mutant mice develop colitis when exposed to opportunistic bacterial pathogens^14^ yet remain disease-free under germ-free conditions^15^. The stark contrast between naturally colonized δD910A mice showing intestinal inflammation and healthy states observed in germ-free settings points to a critical link between pathogen surveillance and immune control mechanisms.

PI3Kδ plays a fundamental role in adaptive immunity, acting downstream of the T-cell receptor (TCR) and B-cell receptor (BCR) to orchestrate key immune functions, including cytokine production, lymphocyte proliferation, and differentiation^16, 17^. PI3Kδ activity is essential for the functions of T helper (Th) cells, CD8⁺ cytotoxic T cells, and B cells, making it a central node in shaping antigen-specific adaptive immunity^16^. Systemic deletion or mutation of PI3Kδ leads to reduced antigen-specific Th cell responses in models such as ovalbumin (OVA)-induced vaccination and myelin oligodendrocyte glycoprotein (MOG 35-55)-induced experimental autoimmune encephalomyelitis (EAE), underscoring its role in antigen-driven T-cell responses^13, 16, 18^. Despite these insights, whole-body PI3Kδ deletion limits the ability to distinguish between innate and adaptive immune contributions to intestinal immune regulation, particularly under inflammatory conditions like colitis, underscoring the importance of employing immune-cell-specific models to resolve the cell-intrinsic roles of PI3Kδ in pathological contexts.

In the dynamic and complex environment of the gut, dendritic cells (DCs) serve as sentinels of mucosal immunity, which, as antigen-presenting cells (APCs), calibrate immunological tolerance towards beneficial dietary antigens and commensal microorganisms while orchestrating robust immune responses against enteric pathogens^19, 20, 21^. Under homeostatic conditions, intestinal DCs take up and transport antigens to mesenteric lymph nodes (mLNs), where they promote tolerance by priming regulatory T cells (Tregs) in the gut^21, 22, 23^. DCs utilize evolutionarily conserved pattern recognition receptors (PRRs) to discriminate between innocuous and microbial entities by detecting pathogen-associated molecular patterns (PAMPs)^24, 25^. Upon PAMP recognition by PRRs, DCs undergo a maturation process, which is characterized by enhanced antigen presentation capacity, increased co-stimulatory molecule expression, and release of inflammatory and immunomodulatory molecules, programming them as potent APCs to instruct specific T-cell responses tailored to precise immunological contexts^26, 27^. The fundamental role of DCs, as APCs, becomes evident when impaired MHC-II-restricted antigen presentation or cDC ablation in vivo leads to tolerance failure, spontaneous autoimmunity, and intestinal inflammation^28, 29^.

Although PI3Kδ functions in adaptive immune cells are well-characterized, a significant knowledge gap remains regarding its specific contributions to DC-mediated regulation of immune responses, particularly in the context of systemic PI3Kδ deficiency, where intestinal inflammation and susceptibility to infections emerge as a prominent pathological feature. Through an integrated experimental strategy combining genetic models, biochemical assays, and immunological analyses, we show that p110δ is essential for coordinating the molecular crosstalk between DC antigen presentation and microbial surveillance capabilities by PRRs, thereby instructing adaptive T cell responses. Our findings reveal a fundamental DC-intrinsic role of PI3Kδ in shaping adaptive immunity and supporting intestinal immune balance. Furthermore, we demonstrate how PI3Kδ deficiency creates conditions conducive to colitis development, revealing that DC-intrinsic immune mechanisms may serve as potential therapeutic targets for restoring intestinal immune homeostasis.

## Results

### Genetic inactivation of p110δ PI3K induces hyperactivation of intestinal DCs preceding enteric bacteria-mediated colitis

Given that enteric bacteria can trigger colitis in δD910A mice, systemically carrying an inactivated p110δ^13, 15^, we hypothesized that PI3Kδ deficiency impairs DC function, compromising microbial surveillance and intestinal immune homeostasis. We further postulated that dysfunctional PI3Kδ-deficient DCs fail to mount effective adaptive immune responses, thus contributing to colitis pathogenesis. To assess our hypothesis, we co-housed δD910A mice with wild-type (WT) mice in conventional (CONV) or specific pathogen-free (SPF) conditions for 35 weeks (Fig. 1a). Shortly after weaning, δD910A mice housed under CONV conditions (δD910A-CONV) exhibited hallmark features of intestinal inflammation, including rectal prolapse, gastrointestinal bleeding, weight loss, and colon shortening, with disease severity peaking between 11 and 15 weeks of age (Fig. 1b–d). Histological analysis revealed marked immune cell infiltration in the lamina propria and epithelial hyperplasia, confirming inflammation and correlating with a significantly elevated disease activity index (DAI) compared to WT-CONV controls (Fig. 1e-f and Supplementary data Table S1). Additionally, Lipocalin-2, a marker of intestinal inflammation^30^, was increased in intestinal mucus secretions from δD910A-CONV mice compared with WT controls (Extended data Fig. 1a). In contrast, δD910A-CONV mice treated with antibiotics or housed under SPF conditions exhibited no macroscopic or microscopic signs of colitis (Fig. 1b, d, and Extended data Fig. 1b).

**Fig. 1:**
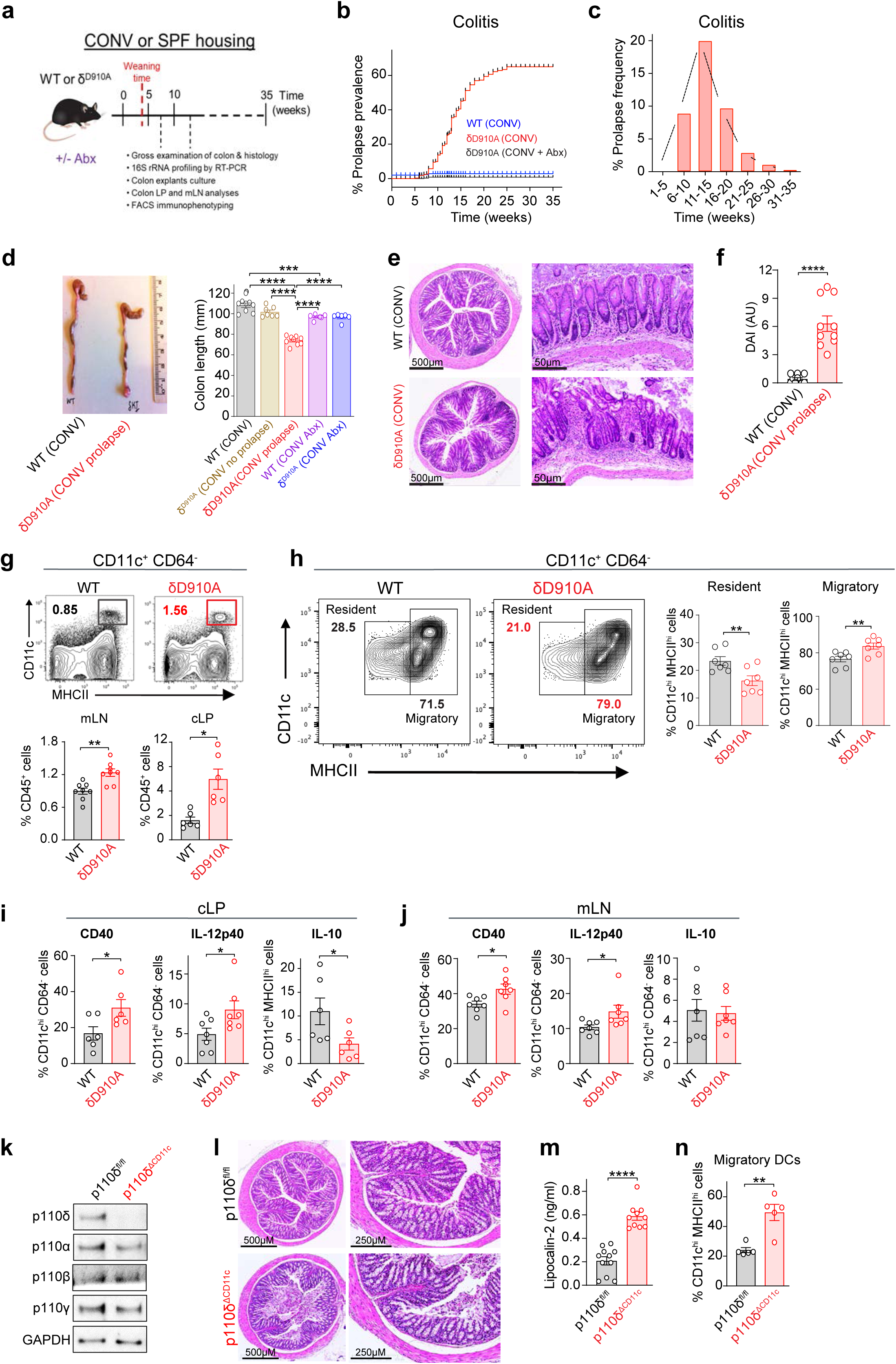
Microbiota-driven colitis overlaps with hyperactive DCs in PI3Kδ deficiency. (**a**) Scheme representing experimental workflow for mice in CONV or SPF housing. (**b**) Kaplan-Meier analysis of age-dependent development of rectal prolapse in WT, δD910A mice (n= 300-356 per group) or WT mice treated with antibiotics (Abx) (n=10-15) in CONV housing. Log-rank (Mantel-Cox) test. ***p<0.0001. (**c**) Prolapse frequency across weeks in δD910A mice (n=275). (**d**) Representative image of colons (*right* panel) and bar charts depicting colon lengths of indicated mouse genotypes, treated with Veh or Abx (*left* panel) in CONV housing (n=5-9 per group). (**e**) Representative images of H&E staining of Swiss roll (left panel) or transversal colon segments (*right panel*) from WT-CONV or δD910A-CONV mice. Scale bars, 500 or 50 µM, respectively. (**f**) Comparative disease activity index (DAI) of WT and δD910A mice in CONV housing (n=10 per group). (**g-h**) cDCs from WT-CONV or δD910A-CONV mice were isolated and analyzed by flow cytometry. (**g**) Representative contour plots (*upper* panel) and proportions (*lower* panel) of cDCs in mLNs and cLP, (**h**) Representative contour plots (*left* panel) and proportions (*right* panel) of resident and migratory cDCs in mLNs (n=7, per group), and (**i-j**) proportions of CD40^+^, IL-12p40^+^, or IL-10^+^ cDCs from (**i**) cLP (n=6-7 per group) and (**j**) mLNs (n=6 per group) are shown. (**k**) Representative immunoblots of class I PI3K isoforms (PI3Kδ, p110α, p110β, and p110γ) in the total cell lysates of splenic DCs (sDCs) from p110δ^fl/fl^ and p110δ^ΔCD11c^ mice. GAPDH was used as loading control (n=3). (**l**) Representative images of H&E-stained Swiss roll (*left* panel) or magnified transversal colon segment (*right* panel) from p110δ^fl/fl^ and p110δ^ΔCD11c^ mice. Scale bars, 500 or 50 µM, respectively. (**m**) Lipocalin-2 levels in colon mucus from indicated mice assessed by ELISA (n=11-12 per group). (**n**) Proportions of migratory cDCs in mLNs of p110δ^fl/fl^ and p110δ^ΔCD11c^ mice, analyzed by flow cytometry. (n=5 per group). Data are shown as means ± SEM, each data point represents an independent biological experiment. One-way ANOVA, two-way ANOVA with Tukey’s post-hoc test, or two-tailed student t-test (unpaired) were performed for statistical analysis, and *p* values were considered as \**p*<0.05, \*\**p*<0.01, \*\*\**p*<0.001, \*\*\**p*<0.0001.

Maladapted host-microbiota interactions in immune-deficient mouse strains promote chronic gut inflammation by favoring opportunistic bacteria that expand in inflammatory conditions^31, 32, 33^. Consistent with the role of enteric bacteria in colitis, δD910A-CONV mice exhibited reduced fecal 16S rRNA levels, altered microbial composition, which coincided with the onset of rectal prolapse (Extended data Fig. 1c). Notably, fecal samples from δD910A-CONV mice showed significant enrichment of *Helicobacter hepaticus* (*H. hepaticus*), a Gram negative bacterium endemic to CONV housing^34^. *H. hepaticus* levels were more elevated in δD910A-CONV mice with subclinical phenotypes compared with WT controls, and bacterial abundance further increased in association with overt colitis pathology. (Extended data Fig. 1d). In contrast, δD910A-SPF mice exhibited no reduction in fecal bacterial biomass compared with WT-SPF controls (data not shown), consistent with the absence of colitis. Collectively, our data suggest that murine PI3Kδ deficiency drives the expansion of opportunistic enteric bacteria, thereby contributing to colitis pathogenesis.

PI3K inactivation in DCs was reported to amplify innate immune response to bacterial PAMPs and increase endotoxin susceptibility *in vivo*^35, 36^. We next assessed the numbers and functional status of DCs in δD910A colon lamina propria (cLP) and mesenteric lymph nodes (mLNs) (Supplementary data Table S2a). Higher proportions of DCs were present in mLNs and cLP of δD910A mice compared with WT counterparts (Figure 1g). A significant increase in DC proportions coincided with the expanded proportions of migratory DCs in mLNs of δD910A mice (Figure 1h). Functionally, DCs in the cLP and mLNs of δD910A mice displayed a hyperactivated phenotype, as demonstrated by an increased proportion of the cells expressing CD40 surface molecule and intracellular IL-12p40, yet in contrast, a lower proportion of cLP DCs expressed IL-10 compared with WT cells (Figures 1i, j). Furthermore, hyperactivated DC phenotype correlated with a pronounced increase in pro-inflammatory cytokine secretion, alongside a reduction in IL-10 levels in colon explants (Extended data Fig. 1d). These data indicate that murine PI3Kδ deficiency causes overgrowth of opportunistic enteric bacteria, leading to colitis, which is accompanied by a hyperactivated DC state in both local intestinal tissue and mLNs.

### DC-selective PI3Kδ deletion disrupts T-cell immune balance and exacerbates DSS-induced intestinal inflammation

To elucidate the DC-specific role of PI3Kδ in intestinal inflammation and colitis pathogenesis, we generated mice with a targeted deletion of the *Pik3cd* gene, encoding p110δ, under the control of the *CD11c-Cre* promoter (hereafter referred to as PI3Kδ^ΔCD11c^). Splenic DCs from PI3Kδ^ΔCD11c^ showed complete and selective loss of PI3Kδ with no alteration in the expression of other class I PI3K isoforms (Fig. 1k). Colon sections revealed detectable tissue inflammation and increased levels of Lipocalin-2 in intestinal mucus secretions of PI3Kδ^ΔCD11c^ mice compared with controls at around 15 weeks onwards (Fig. 1l, m).

To assess how CD11c-specific PI3Kδ deletion influences immune reactivity to enteric components, we examined intestinal homeostasis in PI3Kδ^ΔCD11c^ mice. An increased proportion of migratory DCs in PI3Kδ^ΔCD11c^ mouse mLNs indicated an active immune response within the intestinal tissue (Fig. 1n). Furthermore, PI3Kδ^ΔCD11c^ mLNs, compared to δD910A mLNs, showed an altered CD4:CD8 T-cell ratio (Extended data Fig. 2a, b), which is associated with dysregulated immune function in inflammatory settings^37, 38^. This altered T-cell ratio coincided with significantly increased proportions of IFNγ and IL-17 expressing proinflammatory CD8^+^ T-cells in the mLNs and cLPs of δD910A and PI3Kδ^ΔCD11c^ mice when compared with their respective controls (Fig. 2a-c). Similarly, mLNs of PI3Kδ^ΔCD11c^ mice, akin to those observed in δD910A mice, showed a significant decrease in the proportions of FOXP3^+^, IL-10^+^, and IL-10^+^IFNγ^+^, IFNγ^+^, and IL-17^+^ CD4^+^ T-cells (Fig. 2d-f and Extended data Fig. 2c), suggesting DC-intrinsic PI3Kδ is necessary for efficient priming of CD4^+^ T cells, notably Tregs, in gut-draining lymph nodes. Notably, in homeostatic conditions, the cLP of PI3Kδ^ΔCD11c^ mice maintained Treg proportions comparable to those in control mice, except for a significant decline in FOXP3 expressing CD4^+^ T cells suggesting other APC subsets may buffer to support Treg homeostasis in the cLP (Extended Fig. 2c). Concurrently, the absence of prolapse pathology and histopathological signs in δD910A-CONV animals crossed onto the *Rag2* deficient (*Rag2^-/-^*) background further substantiated the role of pathogenic T-cell responses in driving colitis (Extended data Fig. 2d-f).

**Fig. 2:**
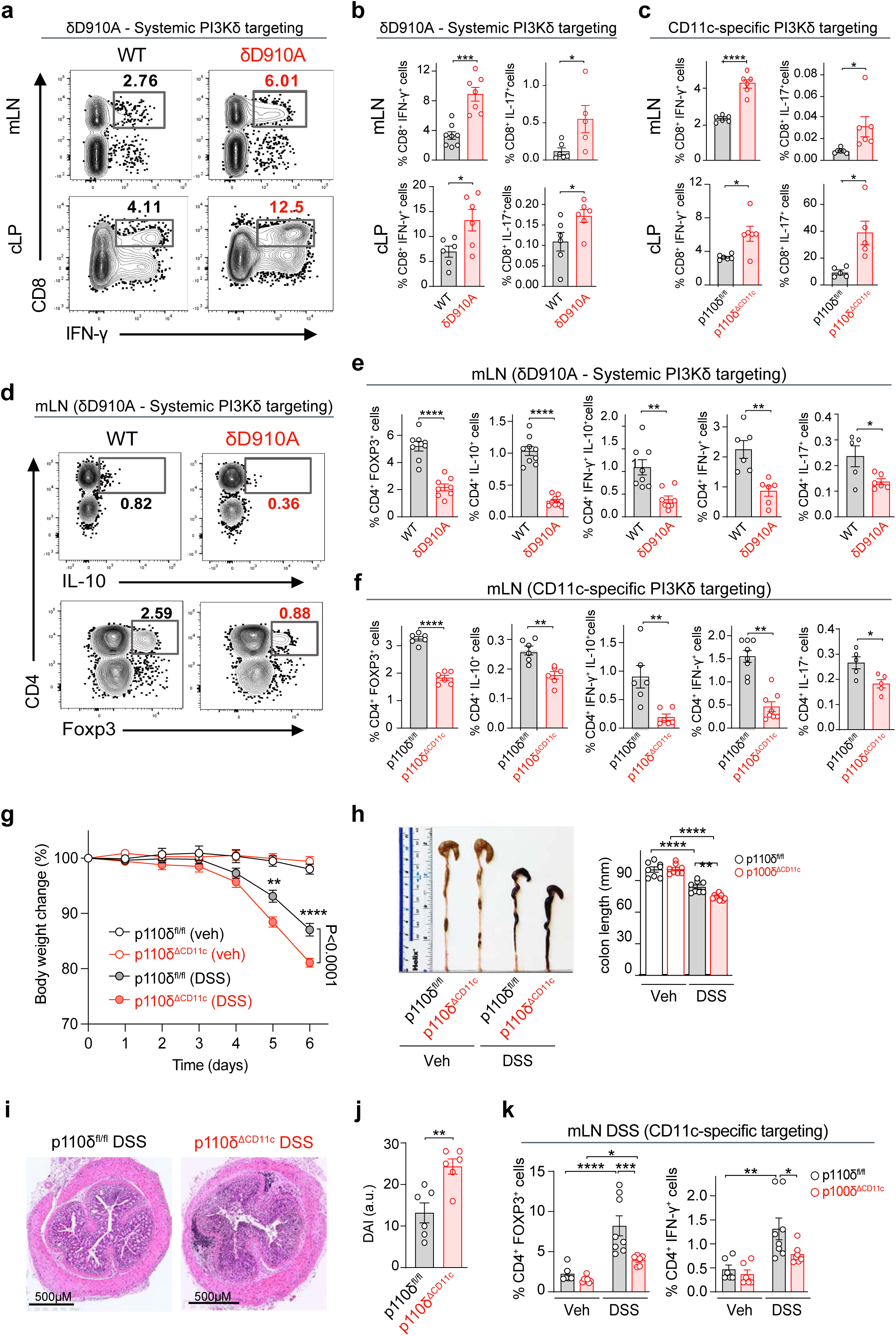
DC-selective PI3Kδ deficiency leads to T-cell immune dysregulation, heightening susceptibility to DSS-mediated intestinal injury. Immune cells from mLNs or cLP of WT, δD910A, p110δ^fl/fl,^ and p110δ^ΔCD11c^ mice were isolated and analyzed by flow cytometry. (**a-c**) Proportions of IFNγ^+^ or IL17^+^ CD8^+^ T cells isolated from (**a, b**) WT and δD910A (**a**) mLNs shown as representative contour plots, or (**b**) mLNs and cLPs as bar charts, or (**c**) p110δ^fl/fl^ and p110δ^ΔCD11c^ mouse mLNs represented as bar charts after PMA and ionomycin stimulation *in vitro* (n=5-7 per group). (**d-f**) Proportions of CD4^+^ T cells in mLNs expressing intracellular (**d**) FOXP3 or IL10 in WT and δD910A mice as representative contour plots, or (**e-f**) FOXP3, IL10, IFNγ, IFNγ IL10, IFNγ or IL17 in (**e**) WT and δD910A, or (**f**) p110δ^fl/fl^ and p110δ^ΔCD11c^ mice as bar charts after stimulation as in *a* (n=5-7 per group). (**g-j**) p110δ^fl/fl^ and p110δ^ΔCD11c^ mice were exposed to DSS or Veh in drinking water for 5 days. (**g**) Body weight loss (N=3, n=4 per group) (**h**) colon length quantification (*left* panel) with a representative image (*right* panel) of indicated mice (n=8-11 per group) (**i**) Representative images of H&E-stained Swiss roll (*left* panel) or magnified transversal colon segment (*right* panel) from p110δ^fl/fl^ and p110δ^ΔCD11c^ mice. Scale bars, 500 or 50 µM, respectively. (**j**) DAI (n=6). (**k**) Proportions of CD4^+^ T cells expressing FOXP3 or IFNγ in mLNs of p110δ^fl/fl^ and p110δ^ΔCD11c^ mice, analyzed after PMA and ionomycin restimulation *in vitro* (n=6-8). Data are expressed as means ± SEM, each data point represents an independent biological experiment. Two-way ANOVA with Tukey post-hoc, one-way ANOVA, or student t-test (unpaired) was performed for statistical analysis, and *p* values were considered as \**p*<0.05, \*\**p*<0.01, \*\*\**p*<0.001, \*\*\**p*<0.0001.

To further investigate how systemic and CD11c-specific PI3Kδ deficiency affects immune responses to enteric microbiota, we first induced acute intestinal barrier injury in mice, housed in CONV or SPF conditions, by oral administration of dextran sodium sulfate (DSS). δD910A mice showed pronounced weight loss, significant shortening of colon lengths, and increased DAI score compared with WT mice in all housing conditions (Extended Fig. 2g-i). Notably, δD910A mice in CONV housing experienced the most severe pathological signs (Extended data Fig. 2g-i). These results suggest that PI3Kδ inactivation amplifies intestinal inflammation in response to commensal microbiota following intestinal injury, with the inflammatory response further intensified by the presence of opportunistic pathogens

In a parallel model of DSS-induced intestinal injury, PI3Kδ^ΔCD11c^ mice developed significantly more severe colitis than the control mice (Fig. 2g-j). The heightened disease severity in PI3Kδ^ΔCD11c^ mice was accompanied by increased monocyte infiltration into the cLP and a marked reduction in FOXP3⁺ regulatory and IFNγ⁺ effector CD4⁺ T-cell populations within the mLNs in the DSS setting (Fig. 2k and Extended Fig. 2j). Overall, CD11c-specific PI3Kδ deletion significantly impaired the expansion of FOXP3 and IFNγ-expressing CD4⁺ T cells following DSS-induced intestinal injury (Fig. 2k). These data indicate that DC-intrinsic PI3Kδ activity is critical for maintaining the balance and expansion of key CD4⁺ T-cell subsets following intestinal injury. In contrast, CD8⁺ T-cell proportions producing IFNγ or IL-17 remained unchanged across genotypes (data not shown), underscoring a selective requirement for PI3Kδ in the CD4⁺ T-cell compartment. Notably, the DC-intrinsic PI3Kδ protective role was further reinforced by the observation that the PI3Kδ^ΔLyZM^ strain, in which PI3Kδ is selectively deleted in macrophage and monocyte populations, exhibited similar levels of intestinal damage and inflammation compared with their controls following DSS exposure (Extended data Fig. 2k, l). Together, these data demonstrate that PI3Kδ activity in DCs is essential for priming protective CD4⁺ T-cell responses, notably the expansion of FOXP3⁺ Tregs, thus ensuring precise immune calibration to microbial cues and injury-associated colonic inflammation.

### Inactivation of DC-intrinsic PI3Kδ dysregulates PRR Signaling and triggers inflammasome activation

Next, we explored whether altered innate responses to conserved PRR pathways, which detect bacterial PAMPs, may underlie the paradox of hyperactivated DCs yet impaired T cell activation in the PI3Kδ-deficient context. Nucleotide oligomerization domain (NOD)1 and NOD2 recognize peptidoglycan-derived dipeptides from bacterial cell wall and play central roles in maintaining intestinal immune homeostasis^39, 40^. NOD2 dysfunction is associated with inflammatory bowel disease (IBD)^41, 42^, and NOD2-deficient (*Nod2^-/-^*) mice have increased susceptibility to *H. hepaticus*-induced colitis^43^. Therefore, we stimulated WT, δD910A or *Nod2^-/-^*bone marrow-derived DCs in the presence of granulocyte-macrophage colony-stimulating factor (GM-CSF) (referred to hereafter BM-DCs) with the NOD1 ligand γ-D-glutamyl-meso-diaminopimelic acid (iE-DAP), NOD2 ligand muramyl dipeptide (MDP), and the TLR4 ligand LPS in WT BM-DCs, NOD2 stimulation with MDP time-dependently increased AKT phosphorylation an established downstream target of class I PI3Ks (Fig. 3a, *upper panel*). AKT phosphorylation was absent in δD910A BM-DCs or when WT cells were pretreated with the PI3Kδ-selective inhibitor IC87114, phenocopying the response in *Nod2^-/-^* BM-DCs, thus confirming that PI3Kδ operates as a downstream effector of NOD2 (Fig. 3a, *middle* and *lower*). In a similar setting, MDP-FITC uptake was comparable between WT and δD910A BM-DCs (Extended data Fig. 3a), confirming that PI3Kδ is not involved in MDP internalization. Likewise, genetic or chemical inactivation of PI3Kδ in BM-DCs inhibited AKT phosphorylation following NOD1 engagement by iE-DAP (Fig. 3b). Furthermore, while NOD2-induced p38 MAPK phosphorylation was increased, no major differences were observed in the NF-κB-associated IKKα/β-phosphorylation in δD910A BM-DCs when compared with WT cells (Fig. 3c).

**Fig. 3:**
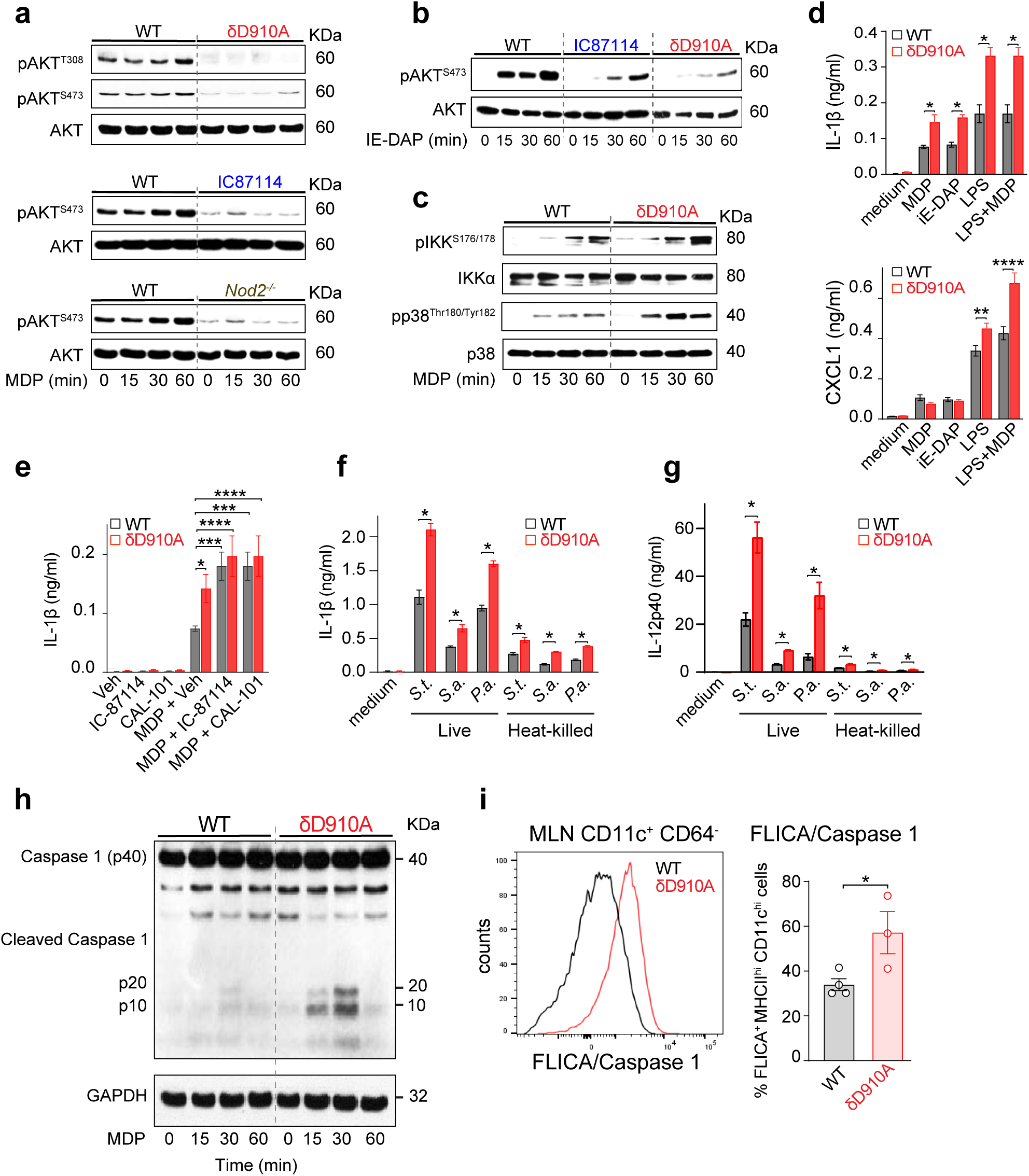
DC intrinsic PI3Kδ fine-tunes PRR signaling mediated by bacterial PAMPs and restrains inflammasome activity. (**a-c)** WT, *Nod2*^-/-^ or δD910A BM-DCs were activated by MDP (NOD2 ligand, 30 µg/ml) or iE-DAP (NOD1 ligand, 30 µg/ml) for the indicated times. (**a, b**) Where shown, WT BM-DCs were pretreated with IC-87114 (PI3Kδ inhibitor) or Veh (DMSO). Representative immunoblots of **(a, b)** p-AKT^Thr308^ and p-AKT^Ser473^, (**c**) p-p38^Thr180/Tyr182^ and pIKK^S176/178^ levels at indicated times are shown, (n=3). (**d**) IL-1β (*upper panel*) and CXCL1 *(lower panel*) levels released by WT or δD910A BM-DCs 18 h post-stimulation with MDP (30 µg/ml), iE-DAP (30 µg/ml), LPS (100 ng/ml) or co-stimulation by MDP+LPS quantified by ELISA (n=3-5 per group). (**e**) Secreted IL-1β levels from WT or δD910A BM-DCs, which were pretreated with IC-87114 or CAL-101 or Veh, followed by MDP stimulation for 18 h (n=3). (**f, g**) WT or δD910A BM-DCs were infected with live or heat-killed (HK) Salmonella Typhimurium (*S.t.*), *Staphylococcus aureus* (*S.a.*), and *Pseudomonas aeruginosa* (*P.a.*) (1:10 ratio) for 15 h. Secreted **(f)** IL-1β and **(g)** IL-12p40 levels in cell culture supernatants were quantified by ELISA (n=5). (**h**) Representative immunoblots of cleaved caspase-1 fragments in WT and δD910A BM-DCs at 90 min after MDP stimulation (n=3). (**i**) Representative flow cytometry histogram (*left*) and percentage of CD11c^+^YVAD^+^ cDCs (*right*) from WT or δD910A DCs from mLNs stimulated with MDP or LPS for 18h (n=3-4). Data are expressed as means ± SEM. Two-way ANOVA with Šídák’s post-hoc test or student t-test (unpaired) was performed for statistical analysis, and *p* values were considered as \**p*<0.05, \*\**p*<0.01, *** *p*<0.001, and **** *p*<0.0001.

Altered PRR signaling in δD910A BM-DCs correlated with increased IL-1β secretion upon stimulation by their cognate ligands; iE-DAP, MDP, and LPS or LPS/MDP costimulation (Fig. 3d, *upper panel*). By contrast, CXCL1 secretion in δD910A BM-DCs was similar to control levels but was significantly higher upon LPS or LPS/MDP costimulation (Fig. 3d, *lower panel*), similar to previous findings^35, 36^. Notably, δD910A BM-DC were phenotypically similar to WT cells, expressing similar surface expression of MHC class II, CD40, and CD86 both at unstimulated and after stimulation by MDP or LPS (Extended data Fig. 3b). Viability assays, analyzed by lactose dehydrogenase (LDH) release and AnnexinV-PI staining, confirmed that cell survival was largely similar between WT and δD910A BM-DCs (Extended data Fig. 3c and data not shown). In WT BM-DCs, treatment with the PI3Kδ-selective inhibitors IC87114 or CAL-101 phenocopied δD910A BM-DC responses, enhancing IL-1β secretion upon stimulation by MDP, iE-DAP, and LPS, compared with vehicle (Veh)-treated WT controls (Fig. 3e). Additionally, δD910A BM-DCs secreted more IL-1β and IL-12p40 after phagocytosis of live or heat-killed (HK) Gram negative S. Typhimurium (*S.t*.), *Pseudomonas aureginosa* (*P.a*) or Gram positive bacterium, *Staphylococcus aureus (S.a.*), with no differences in internalization of live bacteria (Fig. 3f, g and Extended data Fig. 3d). BM-DCs across genotypes exhibited comparable capcity to phagocytose,, confirmed additionally by FITC-coupled HK-*E.coli* uptake, although live *S.t.* uptake showed a slight increase, unrelated to phagocytosis (Extended data Fig. 3d, e).

Hyperactivated DCs have been shown to secrete elevated IL-1β due to inflammasome activity and uniquely remain viable post-activation, enabling superior CD8⁺ T-cell priming.^44, 45,46^. Consistent with increased IL-1β secretion, while δD910A BM-DCs exposure to bacterial PAMPs resulted in pro-caspase-1 processing, WT BM-DCs exhibited little to no detectable pro-caspase-1 cleavage (Fig. 3h). Likewise, δD910A BM-DCs showed substantial pro-caspase-1 processing within 30 min following LPS activation in the absence or presence of extracellular ATP, compared with WT cells (Extended data Fig. 3f, g). Moreover, δD910A DCs from mLNs showed a hyperactivated phenotype with increased levels of active caspase-1 following LPS stimulation *ex vivo*, detected by flow cytometry, using FAM-FLICA probe (Fig. 3i). Additionally, we ruled out mitochondrial ROS (mtROS) dysregulation as a driver of inflammasome hyperactivity in δD910A BM-DCs, since mtROS levels remained unchanged between genotypes at rest or after PRR activation (Extended data Fig. 3h). These findings indicate that PI3Kδ deficiency amplifies bacterial PRR-induced inflammatory responses by enhancing p38 MAPK phosphorylation and inflammasome-mediated IL-1β secretion.

### PI3Kδ deficiency impairs MHC class I and class II restricted antigen presentation in DCs

Our data showed that DC-selective PI3Kδ deficiency dysregulates adaptive T-cell immunity, marked by the pronounced impairment in CD4⁺ T-cell responses despite the hyperactivated phenotype of DCs when activated by bacterial PAMPs. This observation prompted us to investigate whether PI3Kδ deficiency compromises the antigen-presenting function of DCs required for effective T-cell priming. Given that DCs internalize exogenous antigens through multiple endocytic pathways, we carried out in vitro DC-T cell co-culture assays, in which the ovalbumin (OVA) antigen was phagocytosed in particulate form, endocytosed as soluble antigen, or as pre-processed peptides, bypassing the need for intracellular antigen processing (Fig. 4a). Additionally, LPS, as a prototypic PAMP, was coated on the antigenic cargo to determine whether PRR-mediated PI3Kδ signaling selectively modulates the presentation of exogenous antigens internalized via distinct uptake pathways.

**Fig. 4:**
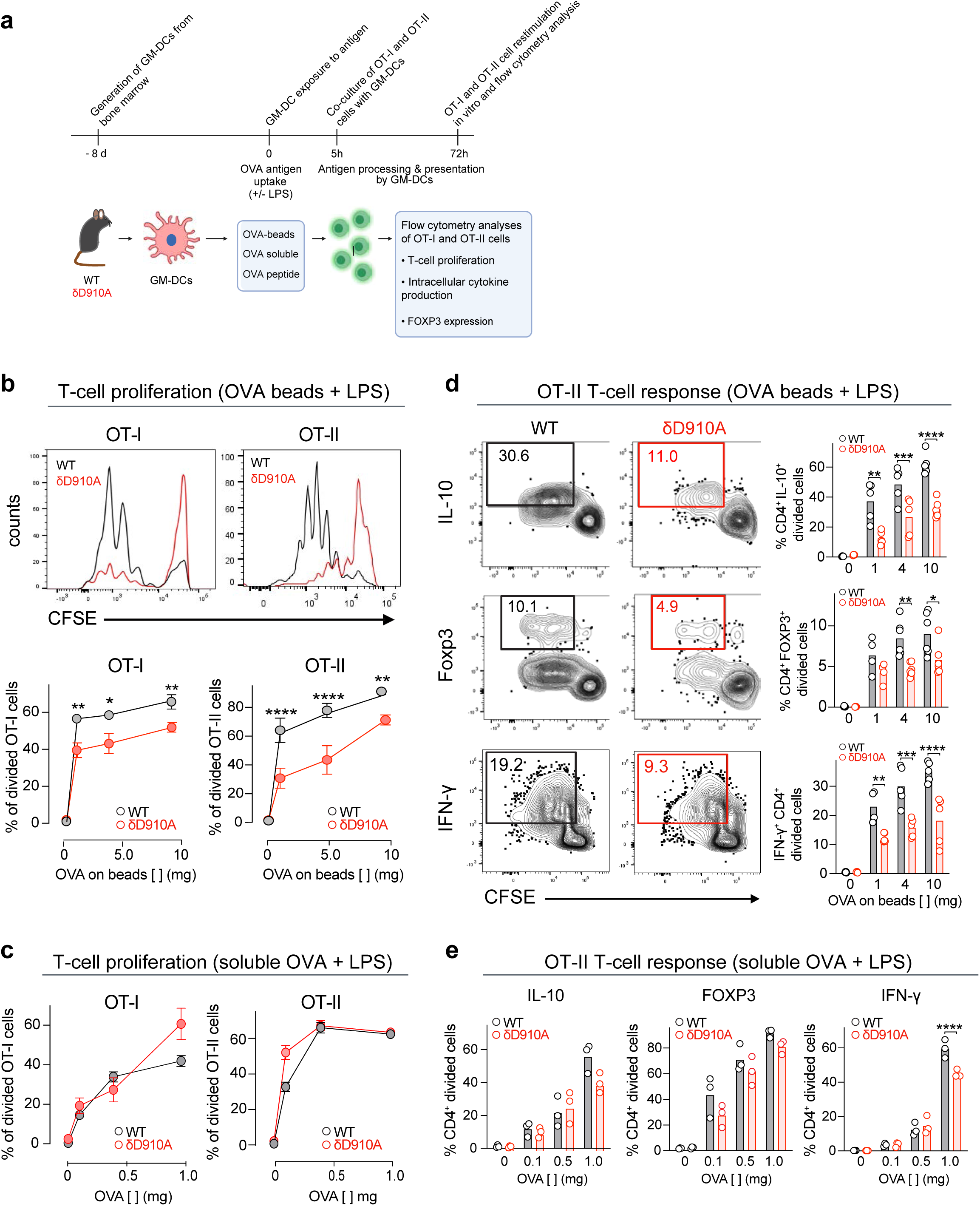
PI3Kδ regulates antigen processing and presentation important for adaptive T-cell responses. (**a**) Schematic diagram showing the antigen presentation assay workflow in BM-DCs. (**b-c**) OT-I or OT-II T-cell division assessed by flow cytometry presented and (**b**) histograms (*upper* panel) and proportions (*bottom* panel) after co-culture with WT or δD910A BM-DCs preloaded with OVA-coated beads (1:50), with graded concentrations of OVA (1-10 mg/ml) in the presence of LPS (100 ng/ml) or (**c**) proportions of divided OT-I or OT-II T cells after co-culture with WT or δD910A BM-DCs, which have endocytosed soluble OVA (0.1-1.0 mg/ml) for 5 h (n=3 per group). (**d**) Representative flow cytometry contour plots (*left* panels*; top, middle,* and *bottom*) and proportions (*right* panels*; top, middle,* and *bottom*) of intracellular IL-10, FOXP3, or IFNγ expressing divided OT-II T cells after co-cultures with indicated genotypes BM-DCs as in *b*, (n=3 per group). (**e**) The proportions of IL-10, FOXP3, and IFNγ intracellular expressing divided OT-II T cells after co-cultures with indicated genotypes of BM-DCs after PMA and ionomycin stimulation *in vitro*, (n=3 per group). Data are expressed as means ± SEM. Two-way ANOVA with Šídák’s post-hoc test was performed for statistical analysis, and *p* values were considered as \**p*<0.05, \*\**p*<0.01, *** *p*<0.001, and **** *p*<0.0001.

Notably, δD910A BM-DCs, compared with WT BM-DCs, showed a significant decrease in the cross-presentation of phagocytosed antigen coated with LPS to OT-I transgenic T cells, which recognize OVA peptide on the MHC class I (Fig. 4b, *left* panels). In a similar setting, δD910A BM-DCs also displayed impaired presentation of phagocytosed OVA coated with LPS to OT-II transgenic T cells, which recognize processed OVA peptides presented on MHC class II (Fig. 4b, *right* panels). In contrast, there was no difference in OT-II T-cell proliferation when BM-DCs from both genotypes presented phagocytosed OVA in the absence of LPS (Extended Fig. 4a), indicating that the requirement for PI3Kδ is contingent on the presence of PAMP. Furthermore, OT-I and OT-II T-cell proliferation remained comparable between WT and δD910A BM-DCs when soluble OVA was used as the antigen, even in the presence of LPS (Fig. 4c). These findings reveal a selective and context-dependent requirement for PI3Kδ in the presentation of phagosome-associated antigens, but not those internalized by endocytosis. Supporting this, direct loading of pre-processed SIINFEKL or OVA323–339 peptides onto BM-DCs resulted in comparable, if not enhanced, T-cell proliferation in δD910A BM-DCs relative to controls (Extended Fig. 4c, d), effectively ruling out intrinsic defects in peptide-MHC complex formation or surface expression. Moreover, core antigen uptake processes, comprising phagocytosis, macropinocytosis, and endocytosis, remained intact as demonstrated by equivalent uptake of latex beads, dextran, or BSA, respectively, across both genotypes in the absence or presence of LPS (Extended Fig. 4e-g and data not shown).

A marked divergence was similarly observed in OT-II T-cell responses, consistent with the T-cell proliferation defects. δD910A BM-DCs induced significantly fewer IL-10^+^, FOXP3^+,^ and IFNγ^+^ OT-II T-cells when compared with WT DCs internalisation of OVA-beads, but not soluble OVA, in the presence of LPS (Fig. 4d). Strikingly, in the absence of LPS and, δD910A BM-DCs elicited comparable, or even elevated, IFNγ expression in OT-II cells when DCs presented phagocytosed OVA antigen (Extended Fig. 4b). These data demonstrate that DC-specific PI3Kδ deficiency selectively impairs MHC class II-restricted priming and cross-priming of T-cells to phagosomal antigens that signal microbial threat.

### PI3Kδ inactivation disrupts NOX2-dependent phagosome oxidative response

To investigate the molecular mechanisms underlying impaired exogenous antigen presentation, we hypothesized that PI3Kδ deficiency impairs NOX2 activity, a key regulator of antigen presentation in DCs. NOX2 generates reactive oxygen species (referred to hereafter as NOX2-ROS), which modulate DC phagosome chemistry, facilitating optimal processing and presentation of exogenous antigens^47, 48^. First, we measured NOX2-ROS production during phagocytosis induced by MDP-coated beads, HK-S.t, and zymosan using luminol-enhanced bioluminescence. WT BM-DCs showed robust and time-dependent increases in luminescence, indicating NOX2-ROS production. NOX2-ROS-mediated bioluminescence was inhibited in WT BM-DCs treated with PI3Kδ-selective inhibitors (PI-3065 or CAL-101) to all stimuli inducing phagocytosis, recapitulating the defect in δD910A BM-DCs (Fig 5a, b and Extended data Fig. 5a). Importantly, the impairment in NOX2-ROS was not due to phagocytosis defects, as uptake of FITC-labeled *E. coli*, latex beads or zymosan was similar between both genotypes (Extended data Fig. 3f, 4e and 5b). The functional significance of impaired NOX2 activity was confirmed in infection assays where δD910A BM-DCs showed a reduced ability to eliminate bacterial pathogens, notably with more *S.t.* and *S.a.* surviving 6 h post-infection (Extended data Fig. 5c). Moreover, no difference in PMA-induced NOX2-ROS generation was observed, which was concurrent with similar expression levels of NOX2 components, p22*phox* or gp91*phox* between both genotypes (Fig. 5c, d). Control experiments using NOX2 subunit gp91*phox* deficient, *Cybb^-/-^* BM-DCs, or WT controls pretreated with antioxidant diphenyleneiodonium (DPI), showed no detectable ROS upon phagocytosis of MDP-coupled beads, HK-*S.t.*, or zymosan (Fig. 5a-d and Extended Fig. 5a).

**Fig. 5:**
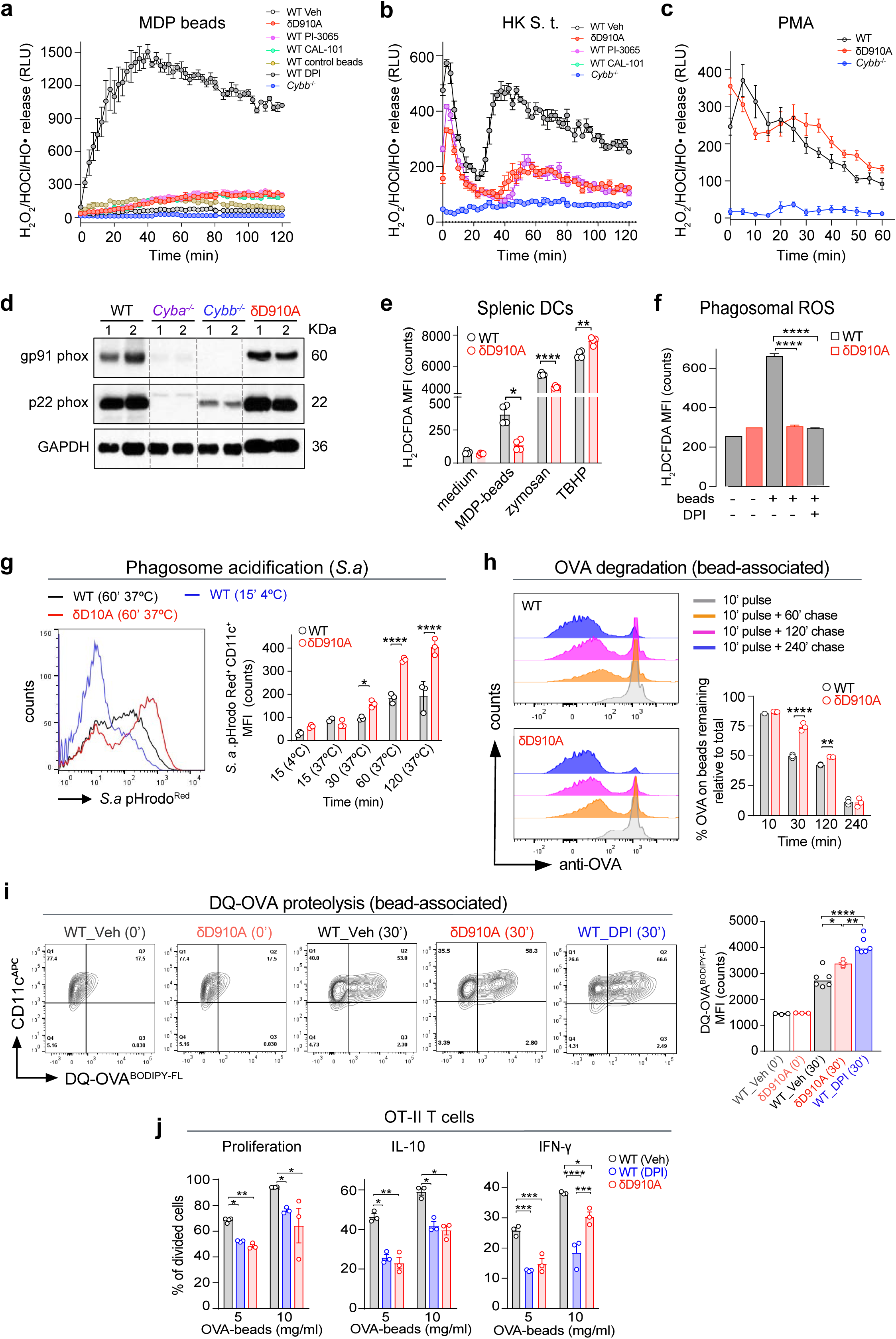
PRR-activated PI3Kδ potentiates NOX2-dependent phagosomal ROS that is essential for an efficient antimicrobial response. (**a, b**) Representative kinetics of total ROS generation from (**a**) δD910A and *Cybb^-/-^*BM-DCs or WT DCs pretreated with PI-3065, CAL-101 or DPI, or Veh that were stimulated with MDP-coated beads or beads alone (3 μm, 1:50), (**b**) δD910A and *Cybb^-/-^* or WT BM-DCs pretreated with PI-3065 or CAL-101 or Veh that were stimulated with heat-killed *Salmonella Typhimurium S.t.* (HK-*S.t.*) or (**c**) WT, δD910A or *Cybb*^-/-^ BM-DCs activated by PMA (1 μg/ml). Luminol-induced chemiluminescence was measured using a Novostar luminometer for up to 2 h. Incubations were performed in triplicates and data (means ± range) from one representative experiment out of 3 are shown and are expressed as relative light units (RLU). (**d**) ROS levels in WT or δD910A splenic DCs (sDCs) were measured as in **e**, at 30 min (n=3 per group). (**e**) Representative immunoblots showing NOX2 components gp91phox and gp22phox in WT, *Cybb*^-/-^, *Cyba*^-/-^ or δD910A BM-DCs total cell extracts. One representative image out of 3 is shown. (**f)** Intra-phagosomal ROS generation measured by flow cytometry following DCFDA-coupled bead phagocytosis in WT BM-DCs, pretreated with Veh or DPI (5 µM), or in δD910A BM-DCs (n=3 per group). (**g**) Representative histogram at 60 min (*left)* and bar chart kinetics (*right*) showing phagosome acidification in WT and δD910A BM-DCs following pHrodo-coupled *S.a.* phagocytosis, assessed by flow cytometry (n=3 per group). (**h**) Representative histogram plots (*left*) and percentage *(right*) of OVA degradation determined by flow cytometry using OVA-specific mAb targeting phagocytosed OVA-coupled beads up to 240 min after chase (n=3 per group). (**i**) Representative counter plots (*left*) and percentage *(right*) of DQ-OVA proteolysis in WT BM-DCs, pretreated with Veh or DPI (5 µM), or in δD910A BM-DCs at 30 min following phagocytosis of DQ-OVA-coupled beads in the presence of LPS (100 ng/ml), determined by flow cytometry (n=3 per group). (**j**) OT-II T-cell proliferation and proportions of divided CD4^+^ T cells expressing intracellular IL-10 and IFNγ, following restimulation by PMA and ionomycin. OT-II T cells were previously co-cultured with DCs previously exposed to OVA-coated beads in the presence of LPS for 5 h. DPI (5 µM) or Veh (DMSO) was added in the indicated groups (n=3 per group). Data are expressed as means ± SEM. One-way ANOVA with Tukey’s post-hoc, or two-way ANOVA with Šídák’s post-hoc test, were performed for statistical analysis, and *p* values were considered as \**p*<0.05, \*\**p*<0.01, *** *p*<0.001, and **** *p*<0.0001.

We next confirmed NOX2-ROS defects in δD910A splenic DCs using the redox-sensitive marker H₂DCFDA (2′,7′-dichlorodihydrofluorescein diacetate), analyzed by flow cytometry. Our data revealed that the ROS defects were specific to phagocytosed cargo containing PAMPs (MDP-coupled beads, HK-S.t, or zymosan) (Fig. 5e). Furthermore, we localized the impaired NOX2-ROS production to BM-DC phagosomes, as demonstrated by the loss of fluorescence in δD910A BM-DCs and DPI-treated WT cells after phagocytosis of DCFDA-coupled beads with LPS (Fig. 5f). Together, these data suggest that PI3Kδ inactivation in DCs impairs NOX2-mediated generation of intra-phagosomal NOX2-ROS and elimination of bacterial pathogens following phagocytosis.

### PI3Kδ regulates phagosomal antigen processing and MHC-II-restricted T-cell priming

Having established the link between PI3Kδ and NOX2 activity, we explored how PI3Kδ inactivation regulates antigen degradation in DC phagosomes and impacts MHC-II-restricted T-cell priming. Previous research suggested that NOX2-ROS can decelerate the acidification rate or overall pH optima in DC phagosomes, thereby creating permissive conditions for the progressive processing of exogenous antigens essential for T-cell priming^47, 48, 49, 50^. Therefore, we assessed acidification kinetics in BM-DC phagosomes, confirming that δD910A BM-DCs acidified more rapidly than WT BM-DCs after internalizing pHrodo-labeled *S. aureus* (Gram-positive), *E. coli* (Gram-negative) or zymosan (Fig. 5g and Extended data Fig. 5d).

Excessive degradation of antigens by lysosomal proteases can impair MHC II-peptide presentation and compromise CD4^+^ T-cell activation, which is counteracted by the action of NOX2-ROS in phagosomes^49, 51, 52^. This led us to explore whether suboptimal antigen processing in δD910A BM-DC phagosomes might explain their impaired T-cell priming capacity. As expected, OVA degradation from phagocytosed OVA-coupled beads progressively increased over time in WT DCs (Fig. 5h). In contrast, δD910A phagosomes exhibited significantly accelerated antigen degradation, with marked differences evident as early as 30 min post-internalization (Fig. 5h). These findings indicate that PI3Kδ is critical for regulating the kinetics of phagosomal proteolysis, and PI3Kδ deficiency leads to premature degradation of antigenic cargo, potentially compromising effective T-cell priming.

To determine whether the rapid loss of OVA in δD910A BM-DC phagosomes was associated with enhanced proteolytic activity due to NOX2-ROS inhibition, we used DQ-OVA^BODIPY-FL^ that emits a fluorogenic signal upon proteolytic cleavage of OVA by proteases in acidic compartments. Soluble DQ-OVA exhibited comparable proteolysis rates between δD910A BM-DCs and WT BM-DCs, with a marginal increase in δD910A cells (Extended data Fig. 5e). By contrast, phagosomal DQ-OVA degradation was significantly elevated in δD910A BM-DCs and was similarly recapitulated in DPI-treated WT cells, though to a greater extent, following internalization of DQ-OVA-coupled beads, indicating increased antigen proteolysis in the absence of PI3Kδ activity (Fig. 5i). Furthermore, δD910A BM-DCs displayed impaired priming of OT-II T-cell proliferation and reduced acquisition of T-cell responses (intracellular IL-10 and IFNγ expression), a defect broadly phenocopied in DPI-pretreated WT BM-DCs presenting phagocytosed OVA antigen coupled to beads in the presence of LPS (Fig. 5j). These findings confirm that PI3Kδ deficiency disrupts NOX2 function, leading to accelerated phagosomal acidification and increased antigen proteolysis, ultimately impairing MHC-II-restricted T-cell priming and immune responses.

#### PRR signaling activates bidirectional crosstalk between PI3Kδ and RAC2

To investigate how PI3Kδ regulates NOX2 activity, we examined its role in modulating RAC2, the predominant RAC isoform in hematopoietic cells^53^, and a known regulator of antigen cross-presentation in DCs^54, 55^. We first assessed the levels of active GTP-bound RAC isoforms (RAC1-3) in WT and δD910A BM-DCs under resting conditions and following stimulation with bacterial PAMPs, MDP, and LPS. While basal RAC activity was similar in both genotypes, only WT BM-DCs exhibited a significant increase in active RAC levels after PRR stimulation (Fig. 6a). Notably, MDP-induced RAC activation was abolished in both δD910A and *Nod2^-/-^*BM-DCs, indicating that NOD2 signaling is required for this acute RAC activation in response to MDP. Pulldown assays confirmed that the MDP-triggered increase in RAC2-GTP was absent in δD910A cells (Fig. 6b), establishing PI3Kδ as an essential potentiator of acute RAC2 activity, downstream of PRR signaling.

**Fig. 6:**
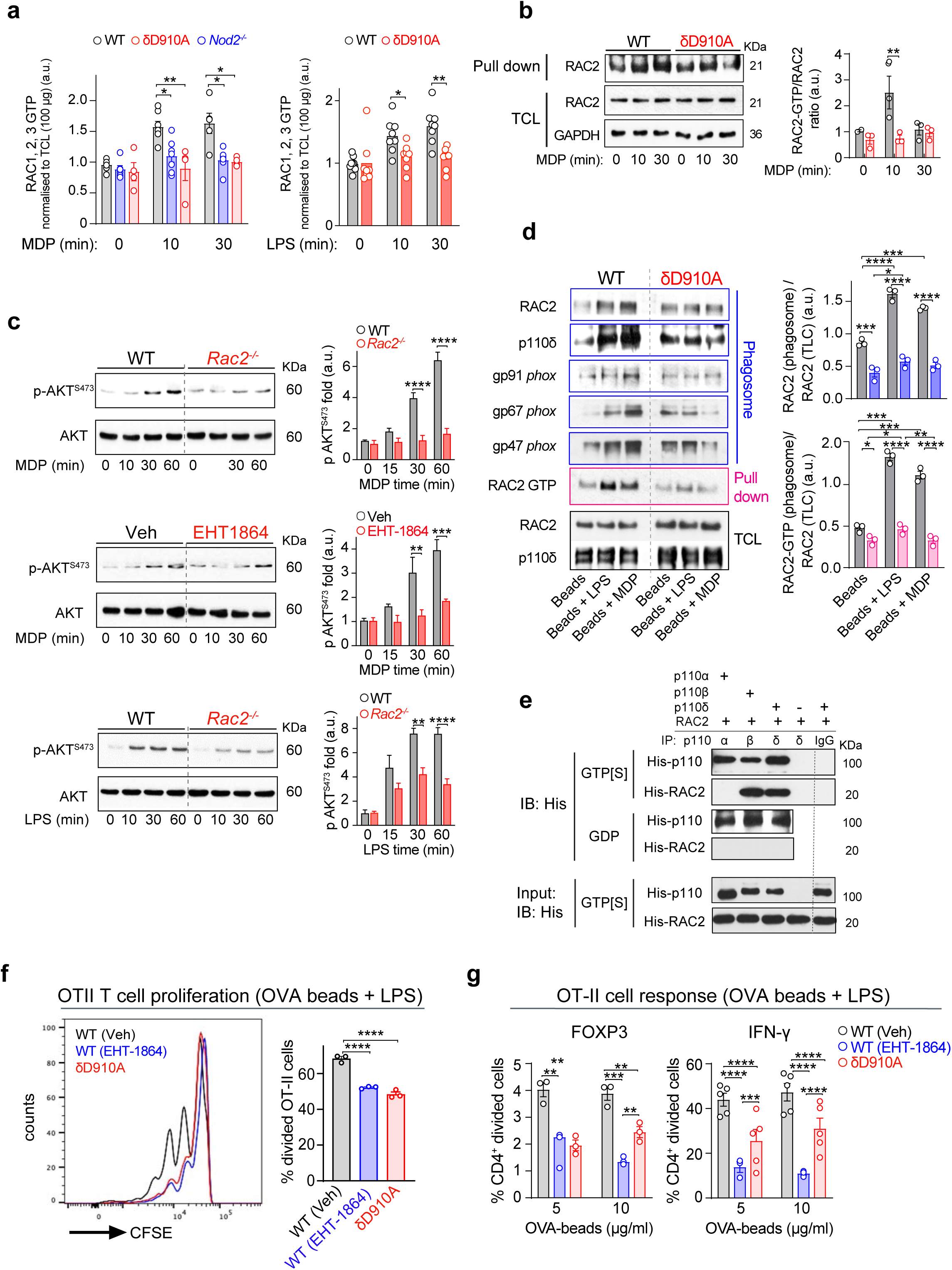
Bidirectional cooperation between PI3Kδ and RAC2 activities on DC phagosomes supports DC antigen presentation function. (**a**) GTP-loaded RAC1/2/3 in WT, δD910A, or *Nod2^-^*^/-^ BM-DCs assessed by Rac G-LISA after stimulation with MDP (30 µg/ml) (*left* panel) or LPS (100 ng/ml) (*right* panel) (n=4-7 per group). (**b**) Representative immunoblots (*left* panel) and bar chart (*right* panel) showing GTP-loaded RAC2 in pulldown complexes and RAC2 in TCL from MDP-stimulated WT and δD910A BM-DCs at indicated time points. GAPDH was used to verify equal loading (n=3). (**c**) Representative immunoblots (*top* and *bottom, left* panels) and fold change (*top* and *bottom*, *right* panels) of p-AKT^Ser473^ levels in WT and RAC2-/- BM-DCs stimulated with MDP (30 µg/ml) or LPS (100 ng/ml) at indicated times. *Middle*, *left,* and *right* panels show WT BM-DCs pretreated for 2 h with EHT1864 (0.5 µM) or Veh (DMSO) before MDP stimulation (n=2-3 per group). (**d**) Representative immunoblots (*left* panels) showing phagosome-associated RAC2, PI3Kδ, and NOX2 components (gp91*phox*, p67*phox*, p47*phox*) in WT or δD910A BM-DCs that phagocytosed 3 µM beads, coated with MDP or LPS, or left uncoated at 30 min. Phagosome-associated RAC2-GTP (magenta rectangle) was pulldown with PAK-CRIB beads from BM-DC phagosome extracts of indicated genotypes, with total RAC2 and PI3Kδ in TCL is shown (black rectangle). Ratios of active RAC2 in phagosomes (*right*, *upper* panel) or total RAC2 (*right*, *lower* panel) are shown (n=3). (**e**) Representative immunoblots showing recombinant His-tagged p110β and p110δ binding to recombinant GTPγS-loaded His-tagged RAC2 in a cell-free system, following immunoprecipitation of complexes using Abs directed against p110α, p110β, or p110δ. GDP-loaded His-tagged RAC2 was used as a control. Immunoblots of complexes were probed using anti-His mAb. Total recombinant protein loading visualized using anti-His mAb. One representative experiment is shown (n=2). (**f**) OT-II T-cell division after co-culture with WT or δD910A BMDCs, pretreated with RAC inhibitor (EHT-1864, 0.5 µM) or Veh (DMSO) before preloading with OVA-coated beads in the presence of LPS for 5 h (n=3 per group). (**g**) Proportions of Foxp3^+^ and IFN-γ^+^ OT-II T cells after restimulation with PMA and ionomycin. Results are expressed as means ± SEM. Statistical analysis by one-way or two-way ANOVA with Tukey’s post hoc test, with *p* values considered \**p*<0.05, \*\**p*<0.01, *** *p*<0.001, and **** *p*<0.0001.

To assess the functional relationship between PRR-activated PI3Kδ and RAC2, we examined the kinetics of AKT phosphorylation in RAC2-deficient (*Rac2^-^*^/-^) BM-DCs following MDP or LPS stimulation. Notably, RAC2 deficiency significantly decreased AKT phosphorylation following stimulation by MDP and LPS. (Fig. 6c). Additionally, p38 MAPK phosphorylation was enhanced in *Rac2*^-/-^ BM-DCs stimulated with MDP or LPS, phenocopying the effects of PI3Kδ inactivation (Extended data Fig. 6a, Fig. 3c, and 5f). The inhibition of AKT phosphorylation was effectively recapitulated in MDP-stimulated WT BM-DCs pretreated with the pan-RAC inhibitor, EHT-1864, confirming that chemical inhibition of RAC mimics genetic ablation of RAC2 in reducing PRR-induced AKT activation (Fig. 6c). Further experiments showed that tyrosine kinase inhibitors (AG-17, AG-213 or Genistein), blocked both AKT phosphorylation and RAC activation in WT BM-DCs following LPS stimulation (Extended data Fig. 6b and 6c), supporting that PRR-induced tyrosine kinases activate PI3Kδ and its target RAC2. These findings uncover an unexpected bidirectional crosstalk between PI3Kδ and RAC2, wherein PRR-mediated activation of PI3Kδ enhances RAC2 activity, while RAC2, in turn, amplifies PI3Kδ signaling.

### PI3Kδ-RAC2 crosstalk on DC phagosomes drives MHC-II-restricted T-cell priming

Next, to define the spatial and functional interplay between PI3Kδ and RAC2, we investigated their activities restricted to the phagosome membranes, where their activities may coordinate signaling events critical for antigen processing and presentation. For these assays, we isolated phagosomes from WT and δD910A BM-DCs after they internalized either PAMP-coated beads or uncoated beads as controls. Analysis of phagosome lysates 30 min post-bead internalization showed distinct protein recruitment patterns for p110δ, RAC2, and NOX2 components (gp91*phox*, p67*phox,* and p47*phox*). Phagosomes containing uncoated beads showed similar protein levels in both WT and δD910A BM-DCs (Fig. 6d). In contrast, PAMP-coated bead phagosomes in WT BM-DCs exhibited significantly elevated levels of p110δ, RAC2, and NOX2 components, while this protein enrichment was diminished or absent in phagosomes from δD910A BM-DCs (Fig. 6d). The RAC2-GTP pulldown assays carried out in phagosome lysates from BM-DCs following internalization of uncoated or PAMP-coated beads (LPS or MDP), confirmed that WT BM-DCs had significantly higher levels of GTP-bound RAC2 on PAMP-coated phagosome membranes compared with δD910A BM-DCs (Fig. 6d).

To investigate whether PI3Kδ and RAC2 directly interact, we performed a cell-free *in vitro* assay to examine potential interactions between recombinant RAC2 and PI3Kα, β or δ isoforms. In co-immunoprecipitation assay, using recombinant, GTPγS-loaded RAC1 and RAC2 proteins, we revealed a strong and specific interaction between RAC2 and p110β/p85α or p110δ/p85α heterodimers (Fig. 6e). Consistent with previous findings^56^, we confirmed that p110β/p85α was the sole heterodimer that interacted with GTP-bound RAC1 (Extended data Fig. 6d). We also validated these interactions in co-immunoprecipitation experiments using NIH-3T3 cells stably expressing WT-p110δ or membrane-targeted and thus constitutively active p110δ-CAAX. Following transient transfection with Flag-tagged RAC2, RAC2 co-immunoprecipitated with p110δ-CAAX, confirming that active p110δ forms a complex with RAC2 on endomembranes in intact cells (Extended Fig. 6e). Overall, these data demonstrate that PI3Kδ enhances RAC2 localization and activity at the phagosome endomembranes, and activated RAC2 in turn amplifies PI3Kδ localization at PAMP-containing phagosomes.

Finally, we examined the functional consequences of impaired PI3Kδ-RAC2 crosstalk. Pretreatment of WT BM-DCs with the RAC inhibitor EHT-1864 recapitulated the impaired ability of δD910A BM-DCs to prime OT-II T-cell proliferation (Fig. 6f). Moreover, OT-II T-cells primed by EHT-1864-pretreated WT BM-DCs, similar to δD910A cells, showed decreased frequencies of FOXP3^+^- and IFNγ^+^-expressing cells among the divided T-cell population (Fig. 6g). These results confirm that PI3Kδ-RAC2 crosstalk is necessary for both antigen-specific CD4^+^ T-cell activation and subsequent T-cell differentiation.

## Discussion

In this study, we have elucidated that PI3Kδ plays a central role in regulating intestinal immune homeostasis by coordinating three key DC functions: microbial detection, modulation of inflammatory signaling, and antigen processing and presentation. Central to our findings, PI3Kδ in DCs integrates PRR signaling with downstream effector pathways through the activation of RAC2 and the NOX2, forming a dynamic signaling axis essential for maintaining phagosomal redox balance, antigen processing, and restraining inflammasome activation (Extended Fig. 6f). DC-intrinsic PI3Kδ deficiency sets off downstream effects with adverse consequences marked by impaired NOX2-mediated oxidative burst, which is essential for sustaining phagosomal pH and ensuring efficient proteolysis of phagocytosed antigens. The resulting redox imbalance triggers inflammasome activation and elevated IL-1β secretion. Furthermore, disrupted pH homeostasis in phagosomes causes accelerated proteolysis of phagocytosed antigens, impairing MHC class I and II restricted antigen presentation, compromising T-cell priming despite a hyperactivated inflammatory phenotype (Extended Figure 6f).

Our study underscores the non-redundant role of PI3Kδ in DCs that support gut immune equilibrium. In vivo deletion of PI3Kδ in DCs disrupts CD4^+^ T-cell expansion under homeostatic conditions, particularly affecting Tregs in the mLNs. This immune disequilibrium was accompanied by skewed T-cell responses in the mLNs and cLP marked by an expansion of pro-inflammatory CD8+ T cells producing IFNγ and IL-17. The gut inflammation under PI3Kδ deficiency shares several key features with previously defined animal models of colitis mediated by the deletion of MHC class II in cDCs^28^ or combined ablation of NOX2 and NOD2, defined by an overt inflammatory response to enteric bacteria in the gut^57^. Under conditions of barrier injury and microbial translocation, such as in the DSS-induced colitis, DC-intrinsic PI3Kδ signaling emerges as a non-redundant pathway necessary for promoting protective CD4⁺ T cell responses to support mucosal immunity and tolerance. While our study focuses on the cell-intrinsic role of PI3Kδ in DCs, we acknowledge the broader immunological context in which DCs interact with multiple cell types, including Tregs and effector T cells, for which the PI3Kδ role is as important beyond the gut^11, 58, 59^. Collectively, our data position PI3Kδ in DCs as a key regulator that supports optimal CD4^+^ T-cell immunity in the gut.

At the molecular level, we identified PI3Kδ as a central coordinator that links microbial surveillance signals to phagosome effector functions of DCs. Our data add further complexity to phagosome maturation beyond PRR-induced inflammatory outputs^60, 61^, by which PI3Kδ activity supports repurposing a conserved antimicrobial effector pathway under NOX2 control, originally developed to neutralize pathogens, for antigen presentation. Overall, the shared phenotypes between PI3Kδ-, RAC2-, and NOX2-deficient DCs point to a PI3Kδ–RAC2–NOX2 axis at the phagosomes that supports their APC function^47, 54, 55^. We found that PI3Kδ spatiotemporally links PRR signals to NOX2 via RAC2 at the phagosomes of unopsonized PAMP-coated cargo, which we simulated using PAMP-coated beads. Similar to our observation with bacterial PAMPs, LPS and MDP, other PRR ligands such as zymosan (via Dectin-1/TLR2) induce NOX2-ROS^62^, and both Dectin1 and TLR2 activate PI3Kδ^35, 63^. Consistent with these studies, PI3Kδ-deficient BM-DCs show a 50% reduction in their capacity to generate an oxidative burst following zymosan uptake. By contrast, FcγR binding of Ig-opsonized antigens was reported to engage PI3Kγ-mediated NOX2-ROS, enabling DC cross-presentation^64^, while both PI3Kγ and δ link N-formyl-methionyl-leucyl-phenylalanine (fMLP)-mediated GPCR signals to NOX2 activity in neutrophils ^65^. These findings underscore isoform-specific PI3K-NOX2 coupling downstream of distinct immune receptors, converging on NOX-dependent oxidative burst, but producing biological outcomes across innate immune cell types.

Importantly, we uncovered a dynamic signaling axis wherein PI3Kδ-driven acute RAC2 potentiation downstream of PRRs occurs through a bidirectional crosstalk, with RAC2 similarly reinforcing PI3Kδ activity. This interdependence was observed in RAC2-deficient BM-DCs that showed approximately 50% reduction in PRR-induced AKT phosphorylation (Figure 6c), in contrast to the hyperactivating RAC2 mutations that increase AKT phosphorylation^66^. We showed that tyrosine kinase inhibitors effectively suppress AKT activation and RAC2-GTP loading, highlighting the essential role of tyrosine kinase signaling. Additionally, our data align with established models in which p110 PI3K isoforms directly interact with RAS and RHO GTPases to amplify the PI3K signaling cascade^5, 56, 67, 68^. In cell-free reconstituted models, class I PI3K net kinase activity is synergistically increased through two distinct but cooperative inputs: the SH2 domain of the PI3K regulatory subunit binding to pY residues on activated RTKs, and the direct interaction of the p110 RBD with activated GTPases^69, 70^.This synergistic mechanism is recapitulated in DCs, where PRR activation triggers a sequential process by PI3Kδ recruitment to PtdIns(4,5)P_2_-enriched cell membranes, followed by the activation and phagosome-associated endomembrane enrichment of RAC2-GTP. Overall, RAC2 activation increases net PI3K signaling through further engagement with PRR-driven signals, creating a regulatory crosstalk that enhances the functional output of both pathways. Our findings suggest a molecular context in which PI3Kδ and RAC2 converge at the phagosomal membrane, optimizing antigen processing and facilitating efficient T-cell priming (Extended Fig. 6e).

An intriguing finding from our study is that PI3Kδ signaling selectively regulates the processing of phagosome-associated antigens of microbial origin, while exerting minimal influence on soluble antigen handling, which involves endocytosis. This selective regulation may suggest an evolutionary adaptation of vesicular trafficking, enabling DCs to tailor their effector functions based on antigen size and microbial threat level. Pathogens, including bacteria and fungi, but also infected cells, require a robust NOX2-mediated oxidative response at the phagosome membranes, thereby engaging distinct antigen processing routes for MHC-restricted priming and cross-priming of T cells. Consistent with this, PAMP co-stimulation with soluble OVA failed to elicit detectable NOX2-ROS in BM-DCs (data not shown). These findings support established models of antigen cross-presentation, in which phagocytosed and soluble antigens undergo divergent intracellular processing routes ^71, 72, 73, 74^.

Compartmentalized NOX2-dependent redox activity during phagocytosis may serve as a critical signal by which DC subsets discriminate between innocuous and harmful antigens. Notably, while cDC1 subsets possess constitutive cross-presentation capacity, cDC2 subsets require PRR priming or inflammatory cues to engage in cross-presentation, a paradigm that aligns well with our data that showing selective coupling of PI3Kδ to the phagosome-associated antigen processing ^73, 75, 76, 77^. Recent studies have emphasized the critical role of NOX2 in fine-tuning antimicrobial defense, restraining inflammasome activity, and the suppression of autoinflammatory/autoimmune reactions^78^. NOX2-ROS modulates antigen processing and epitope selection for the MHC class II presentation, which is particularly important for Treg priming and thus restraining colitis^79^ ^48, 49^.

In conclusion, our study uncovers a critical PI3Kδ-RAC2-NOX2 regulatory axis in DCs that governs antigen presentation and immune tolerance, with significant clinical implications. Disruption of this axis may contribute to an umbrella of primary immunodeficiencies, which manifest as chronic infections, inflammatory diseases or autoimmune susceptibility^53, 80, 81, 82,83^. Our findings suggest that therapeutic strategies aimed at restoring PI3Kδ, RAC2, or NOX2 function or targeting disrupted phagosomal pH balance in DCs hold promise for correcting immune dysfunction. These findings are particularly relevant to PI3Kδ inhibitors, which are clinically effective in chemotherapy and immunotherapy, but are associated with serious adverse effects, including intestinal inflammation and infections^84^. Importantly, the central role of the PI3Kδ-RAC2-NOX2 axis in DC antigen presentation and T-cell activation suggests that selective and temporal modulation of PI3Kδ in DCs could enhance vaccine responses, improve graft outcomes, and refine immunotherapy while minimizing adverse effects. Together, our study provides a conceptual framework for developing safer, more precise immunotherapies and improving clinical outcomes in autoinflammation, infections, and immune-related complications.

## Author contributions

E.A. conceived the project, obtained funding, and designed and supervised the study. M.G.N., L.R.C.V., O.H., R.C.M.C.V., K.K., B.M., and E.A. designed and performed experiments and analyzed the results; J.G.M. and O.H. performed flow cytometry experiments, and analyzed data; A.H. designed and performed *in vitro* bacterial infection studies and analyzed results; A.B., R.S, and L.M.G. performed experiments and analyzed the results; A.F., H.E.C.W., M.P., and B.M. provided key reagents and technical assistance, and B.V. provided early funding for histopathology studies in SPF facility); D.D. performed histopathology analysis and provided technical assistance on colitis studies; and E.A. wrote the paper together with M.G. L.R.C.V, and A. H., D.D, O.H and, B.M. edited the paper.

## Acknowledgements

This work was supported by the grants Medical Research Council (MR/M023230/1) and PCOFUND-GA-2013-60’8765 (E.A.), Barts Charity No. MGU0488 (E.A. and K.K.) and Royal Society No. IES\R2\212104 (E.A. and B.M.) and Arthritis Research UK 19867 (E.A. and B.V.).

B.M. was supported by Institut National de la Santé et de la Recherche Médicale (INSERM), and OH was supported by Great Ormond Street Hospital Children’s Charity (registered charity no.1160024), and B.V. (Cancer Research UK - C23338/A10200). European Union Horizon 2020 research and innovation program under the Marie Skłodowska Curie grant agreements No. 753567 (M.G.N.) and 845908 and (L.R.C.V.). We thank Federica Marelli-Berg (QMUL London) and Brigitta Stockinger (The Crick Institute, London) for their insightful comments on the manuscript, Philip Hawkins (Babraham Institute, Cambridge) and Maria Paula Longhi (WHRI, QMUL London) for reagents and technical advice.

## Ethics declarations

B.V. is a consultant for Pharming (Leiden, The Netherlands) and iOnctura (Geneva, Switzerland) and a shareholder of Open Orphan (Dublin, Ireland). The other authors declare no competing interests.

## Materials and Methods

### Mice

All mice, used in the present study, were on C57BL/6 mice background, bred and fed *ad libitum* in conventional (CONV) (Queen Mary University of London, London, UK) or (when indicated) in specific pathogen-free (SPF) (Charles River, Harlow, UK) animal facilities. WT (C57BL/6), δD910A^13^ (Bart Vanhaesebroeck, UCL Cancer institute, London), p110δ^flox/flox^ mice^85^ (Martin Turner, Babraham Institute, Cambridge) with B6.129P2-Lyz2tm1(cre)Ifo/J)^86^ (referred to as LyzM-Cre, mainly targeting macrophages/monocytes), *Cybb^-/-^* (*Cybb^tm1Din/J^*)^87^, and *Cyba*^-/-88^(*Cyba^tm1a^*, Louise van der Weyden, Welcome Trust Sanger Institute, Cambridge), *Rac2^-/-89^* (Heidi Welch, Babraham Institute, Cambridge) and OT-I B6.SJL.CD45.1^90^ and OT-II B6.CD45.2^91^ (referred to as OT-I and OT-II mice) and Tg(Itgax-cre)1v^92^ (referred to as CD11c-Cre, mainly targeting DCs) - Caetano Reis e Sousa, The Francis Crick Institute, London) were generously provided for use in the study. p110δ^flox/flox^ mice were crossbred with CD11c-Cre or LysM-Cre to ablate *Pik3cd* gene in DCs or macrophages/monocytes, respectively, to generate p110δ^ΔCD11c^ or p110δ^ΔLysM^ in the CONV facility. Unless otherwise noted, both sexes were used in all experiments. All animal procedures were conducted according to the requirements and with the approval of the UK Home Office Animals (Scientific Procedures) Acts, 1986. Animal experimentation was approved by the Queen Mary University of London research ethics committee and was carried out under the Home Office license (PPL70/7447).

### Reagents cells and antibodies

All reagents are listed in Table S3. *Salmonella Typhimurium*, *S. aureus*, and *P. aeruginosa* were used and were gifted by David Holden (Imperial College, London) and Abderrahman Hachani (University of Melbourne, Melbourne). All inhibitors or Veh (DMSO ‒ dimethyl sulfoxide) were added between 1-4 h before cell stimulations. NIH-3T3 cell line (American Type Culture Collection) used in the generation of cell clones stably expressing 5′ Myc-tagged human p110δ with or without the CAAX domain in the pMX vector under a neomycin selection cassette were described^35^.

### Histology and colitis scoring

Mice were evaluated for clinical signs and gross indicators of pathology. For histopathological analysis, sections from the proximal, medial, and distal colon were fixed in 4% paraformaldehyde for 18 h. Tissue embedding, sectioning, and staining for hematoxylin and eosin was carried out by Propath, UK. The sections were assessed and scored using a colitis scoring protocol (Table S1) as previously described^93^ by an expert pathologist in a double-blinded manner. The calculated score was presented as the average of individual scores from the indicated sections of each sample analyzed.

### DSS colitis model

Age (10–12-week-old), sex- and housing-matched mice from CONV or SPF facilities were used for all experiments. Colitis was chemically induced by oral administration of 2.5% w/v solution of DSS (MP Biomedicals) in the drinking water over five consecutive days, starting from day 0. Control groups received the same amount of drinking water. Mice were checked daily for general health parameters, weight fluctuations, and clinical indicators of inflammation, for 7 days. A 20% weight reduction from baseline was predetermined as the humane endpoint for terminating animal participation in the experiment. The DAI (Disease Activity Index) calculations included weight loss, stool consistency, and rectal bleeding were used to assess disease activity as described in Table S3 and were previously described^93^. Body weight loss was calculated as the percentage difference between the original body weight on day 0 and the actual body weight on any day and animals.

### Isolation of cells from cLP and mLNs

Colons were isolated and digested as previously described^94^. Briefly, colons were cut longitudinally, washed with HBSS without Mg^2+^ or Ca^2+^ (ThermoFisher Scientific) and then were trimmed into 1 cm pieces. The epithelial cells were removed by 1.0 mM EDTA (ThermoFisher) of 20 ml HBSS, stirring at 250 rpm for 30 min at 37°C. The colon pieces were washed twice with 10% FBS containing RPMI. Tissues were sliced and processed in 10 ml of a Digestion solution containing 10% FBS RPMI, 200 μg/ml DNase I (Merck SA), 2 mg/ml Collagenase IV (Roche Products Ltd.) and Dispase II (Merck SA) by stirring at 250 rpm for 30 min at 37 °C. Followingly, digested tissue was filtered through a 70μm cell strainer. The collected cells from the cLP were washed twice in 10% FCS RPMI, counted, and resuspended in appropriate volume for further assessment. The mLNs were placed in a digestion solution for 30 min at 37 ° C at 250 rpm. The resulting material was washed with 10% FBS RPMI, filtered with a 70μm cell strainer, counted, and resuspended in appropriate volume for further assessment.

### Isolation of splenic DCs

DCs and splenocytes were isolated from mouse spleens by digestion for 30 min at 37 °C with 2 mg/ml Collagenase IV and DNase I (both from Roche). Samples were treated for 2 min at 21 °C with RBC lysis buffer (eBioscience) and washed twice with Hank’s balanced salt solution. Cells were filtered in a 70 μm cell strainer, counted, and resuspended in appropriate volume for further assessment. When required, splenic CD11c^+^ cells were purified by positive selection with magnetic CD11c^+^ beads according to the manufacturer’s instructions (Miltenyi Biotec.).

### Generation of BM-DCs

BM-DCs were generated from mouse bone marrow precursors as described previously^95^. Cells were cultured for 8-10 days in RPMI-1640 medium (Invitrogen) supplemented with glutamine (Invitrogen), penicillin (Invitrogen), streptomycin (Invitrogen), and 10% heat-inactivated FBS (Invitrogen) in the presence of 20 ng/ml recombinant murine granulocyte-macrophage colony-stimulating factor (GM-CSF) (PreproTech). The viability and absolute number of BM-DCs were assessed by trypan blue exclusion before assays. The cells obtained were routinely >90% CD11c^hi^ by flow cytometry.

### Flow cytometry

Single-cell suspensions were incubated on ice with conjugated antibodies in PBS containing 0.5% BSA and 1 mM EDTA. Unlabeled anti-CD16/32 (clone 2.4G2, BD Biosciences) was used to block Fc receptors together with LIVE/DEAD™ Fixable Aqua Dead Cell Stain (Thermofisher) on ice-cold PBS for 15 min. Next, the cells were washed and incubated with selected antibodies (Table S3) in PBS buffer containing 0.5% BSA for 30 min *at 4°C*, washed two times, and assessed by Flow cytometry. For intracellular staining, cells were fixed and permeabilized using the FOXP3/Transcription Factor Staining Buffer Set according to the manufacturer’s instructions (Thermo Fisher Scientific). Briefly, cells were treated with the FOXP3 Fixation/Permeabilization working solution for 1 h on ice, followed by intracellular staining with conjugated antibodies (Table S3) in 1X permeabilization buffer for 30 min on ice. For intracellular cytokine staining, T cells were pre-stimulated at 37°C for 4 h in RPMI medium supplemented with 10% FBS, phorbol 12-myristate 13-acetate (PMA) (20 ng/ml), ionomycin (4µg/ml), and protein transport inhibitor (ThermoFisher). Intracellular staining of mLN or cLP DCs, cells were stimulated by LPS (100 ng/ml) or left in the medium at 37°C for 6 h. The caspase-1 activity in MLN DCs was assessed using Green Fluorescent FAM-FLICA® caspase-1 (FAM-YVAD-FMK) probe (2BScientific Ltd.) according to the manufacturer’s instructions. Flow cytometry analyses were performed on BD LSRFortessa™ Cell Analyzer (Wokingham, UK). Data were acquired by Diva software and analyzed using FlowJo software (Oregon, USA).

### Isolation of fecal RNA, real-time RT-PCR

RNA was isolated from stools using the PowerViral® Environmental RNA/DNA Isolation Kit (QIAGEN). SuperScript VILO cDNA synthesis Kit (ThermoFisher Scientific) was used to generate cDNA. Reverse-transcription and real-time PCR were done with CFX Connect Real-Time System of BIO-RAD, using primers to quantify mRNA for universal 16S rRNA or *Helicobacter hepaticus* 16S rRNA (Table S3). The abundance of each specific mRNA was normalized to that of 16S levels in the same sample using the 2^−ΔCt^ method and expressed as relative stimulation.

### Infection and enumeration of intracellular bacteria

Infection of BM-DCs was performed as described^96^. Briefly, log-phase *S. enterica* serovar typhimurium NCTC 12023 (gift from David Holden, Imperial College, London) or *S. aureus* (*Staphylococcus aureus subsp. aureus strain Wood 46*, ATCC) strains were washed with ice-cold PBS, followed by adjustment of bacterial cell number to an OD_600_ of 0.01 (equivalent to 1.0 x 10^7^ cells/ml). The multiplicity of infection (MOI) was adjusted to 10. BM-DCs and bacteria were mixed, and bacteria were allowed to bind to BM-DCs on ice. Infection was initiated by moving the BM-DC-bacteria mixture to a 37°C incubator with 5% CO2. After 30 min of infection, the host-pathogen mixture was washed three times with warm PBS to remove non-adherent BM-DCs and excess extracellular bacteria, which were further eliminated by using 100 g/ml gentamicin for 1 h. BM-DCs with internalized bacteria were placed in complete RPMI without antibiotics at 37oC for 2 h to assess bacteria uptake or 6 h to examine bacterial growth. At indicated times, BM-DCs were lysed in PBS containing 0.1% Triton X-100 to release intracellular bacteria, followed by quantification of the live bacteria through serial dilution after plated on Luria agar or trypticase soy agar plates, counting the CFUs at 18 h.

### DQ-OVA degradation assay

BM-DCs (5 x 10^5^ cells per time point) were pulsed with 1µg/ml DQ-OVA (ThermoFisher) at 37°C for 15 min and then incubated at 37°C for indicated time intervals. After chasing, cells were placed on ice to avoid further degradation and immediately analyzed by flow cytometry. For bead-associated DQ-OVA degradation assay, 3µm latex beads were coated with 100 µg/ml DQ-OVA in PBS at 4°C under rotation overnight. After three washes with ice-cold PBS, OVA-DQ coated beads (1:10) were used to pulse BM-DCs (10^6^ cells per time point) at 37 °C for 15 min. Cells were chased at 37 °C for indicated time points, placed immediately on ice to avoid further protein degradation, and then analyzed by flow cytometry.

#### Bead-coupled OVA degradation analyses

Phagosomal OVA degradation was assessed as previously described^47^. Briefly, 3µm aliphatic amine latex beads (Invitrogen) were covalently coated with OVA (EndoFit, InvivoGen) overnight at 4°C under rotation. Beads were washed 3 times with ice-cold PBS to remove the unbound OVA. 1 x 10^6^ BM-DCs per time point were pulsed for 15 min at 37°C and chased with OVA-coated beads (0.5-10 mg/ml) at indicated time points at 37°C. After chasing, cells were lysed in 50 mM Tris-HCl (pH 7.4) supplemented with 1 mM DTT, 0.5% NP-40, 150 mM NaCl, and protease inhibitor cocktail (Roche). The beads were stained with a rabbit polyclonal anti-OVA (Merck) and a secondary goat anti-rabbit Alexa 488 (Thermofisher).

#### Measurement of DC oxidative burst and detection of total ROS

BM-DCs (1 x 10^6^/ml) or splenic DCs (0.5 x 10^6^/ml) were incubated in DMEM without Phenol Red (Gibco), supplemented as above at 37°C. BM-DCs were pre-treated with p110δ inhibitors (1 µM IC87114, 0.5 µM CAL-101 or Veh (DMSO)) for 1 h before stimulation by PMA (1μg/ml), zymosan (5μg/ml, InvivoGen) or 3 μm Sphero^TM^ polystyrene beads (at a ratio of 1:50, Spherotec, Chicago USA), which were previously coated with MDP (30 μg/ml) or LPS (100 ng/ml). The kinetic measurements of the oxidative burst in cells were obtained using luminol-dependent chemiluminescence (50 μM final concentration, Sigma). Light emission was recorded by the Novostar plate reader (BMG Labtech, Germany) at 37 °C for 120 min. In other experiments, 1 × 10^5^ cells were incubated in DCFDA/H_2_DCFDA (20 µM final concentration, Abcam) for 30 min at 37°C in the dark, washed and resuspended in RPMI-1640 containing no phenol red (Gibco) in a black, clear-bottom 96-well microplate with 100,000 stained cells per well and treated with the ligands (indicated above). Fluorescence was measured on a plate reader at Ex/Em = 485/535 nm in end-point mode on Novostar plate reader (BMG Labtech, Germany) at 37 °C every 30 min for 1 - 2 h. For phagosomal ROS measurement, 3 µm beads were covalently coupled with 20 mM DCFDA (Abcam) in sodium hydrogen carbonate buffer (pH 8) rotating at 4°C overnight as previously described^47^. The beads were washed 3X with cold PBS to remove the unconjugated DCFDA. BM-DCs (10^6^ cells per time point) were incubated for 30 min at 37°C with DCFDA-coated beads (1:10). Phagosomal ROS was detected by flow cytometry using BD FACsaria.

### *In vitro* antigen presentation assays

For DC:T cell co-culture experiments, 8 × 10^4^ BM-DCs were incubated with OVA or BSA in the soluble form (graded OVA concentrations: 0.1, 0.4 and 1 mg/ml) or particulate form (1, 4, and 10 mg/ml), which were coupled on to 3µm beads (see section *bead-coupled OVA degradation*) in presence of LPS (100 ng/ml) for 5 h at 37°C. Next, the BM-DCs were washed three times in complete RPMI (see BM-DC generation). OT-I or OT-II cells were labeled with CFSE (5 µM) (ThermoFisher) in 5 mL PBS at 37°C for 10 min, washed three times in complete RPMI and were then added to the DC co-culture in 96-well round-bottom plates at a ratio of 1:10. After 72 h, at 37°C incubator (5% CO2), OT-I or OT-II cells were restimulated in vitro for 4 h at 37°C with PMA (20 ng/ml), ionomycin (4µg/ml) and protein transport inhibitor (ThermoFisher). For intracellular cytokine expression, T cells were stained with the indicated Abs (Table S3) and analyzed by flow cytometry. For processed peptide presentation, BM-DCs were incubated at 37°C for 1 h with OVA 257-264 (SIINFEKL) or OVA 323-339 (0.1, 0.4, 0.7, and 1.0 µg/ml, all purchased from InvivoGen) peptides.

#### Phagocytosis assays

2 × 10^5^ BM-DCs were plated in 94-well plates and incubated with indicated reagents at 37°C. For latex bead uptake analysis, BM-DCs were incubated with 3µm beads coupled to tetramethyl rhodamine (1:10) (ThermoFisher). BM-DC uptake of *E. coli* (1:100 *E. coli* (K-12 strain) BioParticles™, Alexa Fluor™ 488), zymosan (2 µg/ml Alexa Fluor 488 conjugated Zymosan A). After incubation, BM-DCs were placed on ice and analyzed by flow cytometry. As binding control, cells were incubated with the reagents on ice for the indicated times.

#### Phagosome acidification

5 × 10^5^ BM-DCs per condition were plated in 96-well plates, pulsed for 15 min at 37°C, and then chased at 37°C for the indicated time intervals. To assess phagosome acidification, BM-DCs were incubated with 2 μg/ml of zymosan-pHrodo, *S. aureus-*pHrodo, or *E. coli-*pHrodo (all from Invitrogen) at 37°C for 1 h. After incubation BM-DCs were placed on ice and analyzed by flow cytometry.

#### Phagosome isolation using magnetic beads

Phagosome isolation was performed as previously described^47^. BM-DCs were pulsed with 3μm magnetic beads (Dynabeads™ M-280 Streptavidin, ThermoFisher) in presence or absence of LPS (100 ng/ml) or MDP (30 μg/ml) at 37°C for 30 min. Cells were disrupted in homogenization buffer (8% sucrose, 3 mM imidazole, 2 mM DTT, 58ug/ml DNase, and EDTA-free protease inhibitor). Phagosomes were isolated using a magnet, washed three times in cold PBS, and lysed in total cell lysis buffer (for immunoblotting) or RAC lysis buffer (see section *Rac pull-down assay*). After 20 min at 4°C, the lysates from phagosomes were isolated and frozen at -80°C for immunoblot or RAC pulldown analysis.

#### Enzyme-linked immunosorbent assay (ELISA)

IL-1β, TNF, IL-10, IL-12p23, IL-12p40, IL-12p70, IFN-γ, and IL-17 from BM-DC (5 × 10^5^/ml) culture supernatants and Lipocalin-2 from colon mucus were detected using specific DuoSet ELISAs (Table S3) following the manufacturer’s instructions (R&D Systems).

#### Immunoblotting and immunoprecipitation assays

After cell stimulation, cells were washed in PBS and lysed for 30 min on ice in the appropriate buffer for the subsequent analysis; for immunoblot analysis, total cell lysis buffer (20 mM Tris-HCl, pH 7.5, 1% IGEPAL, 10% Glycerol, 150 mM NaCl, 10 mM NaF, 10 mM Na_3_VO_4_ and 10 mM beta-glycerophosphate); or for immunoprecipitation (IP) buffer (50 mM Tris, pH 7.4, 1% IGEPAL, 10% Glycerol, 5 mM MgCl_2_, 100 mM NaCl, 1 mM DTT, 10 mM Na_3_VO_4_, 10 mM NaF and 10 mM beta-glycerophosphate), supplemented with a cocktail of protease and phosphatase inhibitors (Roche). Protein concentration in total cell lysates and phagosome lysates was measured using the Pierce BCA Protein Assay Kit. 50 ug of total cell lysate were mixed with Laemmli buffer and separated on 10-12% SDS-PAGE, transferred to nitrocellulose membranes, and immunoblotted. Abs used for immunoblotting and immunoprecipitation analysis are listed in Table S3.

#### Rac1,2,3 G-LISA Activation Assay

Rac G-Lisa was performed per the manufacturer’s instructions (Cytoskeleton, Inc.). Briefly, BM-DCs were seeded at 1 × 10^6^/ml per condition, where indicated, cells were preincubated with AG213 (), stimulated with indicated PRR ligands, or left in medium. Cells were lysed using *RAC lysis buffer* containing 50 mM Tris pH 7.4, 10 mM MgCl_2_, 300 mM NaCl, 2% IGEPAL, 10 mM beta-glycerophosphate, and protease inhibitors. Next, 50µg of each TLC was fractioned onto the 96-well Rac-GTP binding plate. Ice-cold *Binding buffer* was added before incubating for 30 min at 4°C in gentle shaking. Then, anti-Rac mAb (recognizing RAC1, 2, and 3) was added to the well for 45 min at RT. The wells were washed three times, and HRP-linked secondary Ab was added for 45 min at RT. After three washes, the *Antigen Presenting buffer* (Cytoskeleton, Inc) was added for 2 min, and Rac-activity was detected using a microplate spectrophotometer at 490 nm.

#### Rac pull-down assay

BM-DCs were seeded at 5 × 10^6^ cells per condition and then stimulated with MDP (30 μg/ml) or LPS (100 ng/ml). Plates were then washed with ice-cold PBS and lysed in modified RAC lysis buffer (50 mM Tris, pH 7.4, 2% IGEPAL, 10% glycerol, 10 mM MgCl_2,_ 100 mM NaCl, 10 mM Na_3_VO_4_, 10 mM NaF, 10 mM beta-glycerophosphate, and protease inhibitors). After centrifugation, an aliquot of TCL in Laemmli buffer was kept, and 100 ug of each lyste was incubated with 25 μl of PAK-GST protein beads (Cytoskeleton, Inc.) in rotation at 4°C for 2 h. Subsequently, beads were extensively washed five times with lysis buffer and resuspended in Laemmli sample buffer for immunoblotting analysis.

#### RAC GTPase loading and recombinant RAC1 and 2 interaction assays

RAC1 or RAC2 GTPase activities were measured using recombinant proteins as previously described^43^. Briefly, recombinant human His-tagged RAC1 or RAC2 proteins (Cytoskeleton, Inc.) were reconstituted at 1 mg/ml in dH_2_O with 1 mM DTT. GTPγS or GDP (Cytoskeleton) was reconstituted to 20 mM with 50 µl of de-ionized water. For GTPγS /GDP loading assays, 10 ng of RAC1 or RAC2 protein was incubated with 2 mM GDP or GTPγS in 50 μl of GTPase loading buffer (20 mM Tris-HCl, pH 7.4, 5 mM EDTA, 25 mM NaCl). After 20 min at 37°C, MgCl_2_ was added to a final concentration of 10 mM on ice. Next, 5 ng of His-tagged p110α-, p110β- or p110δ-85 fusion proteins (Abcam) were incubated with recombinant RAC proteins diluted in binding buffer (20 mM Tris-HCl, pH 7.4, 150 mM NaCl, 5 mM MgCl_2_, 1% Triton X-100, 5 mg/ml BSA) at a final volume of 300 µl. Finally, 2 µg/ml of anti-p110α, β, or δ Abs (see Table S3) was incubated in each condition, except for the control anti-rabbit IgG mAb. The samples were rotated at 4°C for 4 h, followed by the addition of 30 µl 1:1 slurry Protein G beads for another 2 h. Beads were washed five times and then diluted in RAC lysis buffer containing Laemmli buffer.

#### NIH-3T3 cell transfection

5 × 10^5^ NIH-3T3 cells were transfected with 2 ug of Flag Rac2 in pcDNA3 (Plasmid #12192, Addgene) together with 2 µg Myc-tagged p110δ-WT or Myc-tagged p110δ-CAAX plasmids or empty vector pMX using jetPRIME^®^ (Polyplus). After 18 h, transfected cells were collected and washed with ice-cold PBS and then were lysed in an immunoprecipitation buffer (described in the immunoblotting section) for 20 min. The TLC was collected, 50 µg of TCL was kept for loading control in Laemmli buffer, and 2 µg of anti-p110δ Ab was added onto 100 µg of TLC rotating 4°C overnight, followed by 30 µl 1:1 slurry of Protein G beads rotating at 4°C for 2 h and washed 5 times. The beads were diluted in total cell lysis buffer containing Laemmli buffer.

### Statistical analysis

Statistical analyses were performed using GraphPad Prism 9. *P* values for datasets were determined by an unpaired two-tailed Student’s t-test with a 95% confidence interval for comparison between two independent groups. For repeated comparison between the two groups, a multiple unpaired two-tailed t-test was applied. For the assessment of more than two groups, one-way analysis of variance (ANOVA) or two-way ANOVA, followed by Tukey’s or Holm Šídák’s multiple comparison tests, was used. Kaplan-Meier survival analysis was used to estimate the incidence of anal prolapse and the non-parametric *log-rank* test for the frequency of prolapse across age of mice.

### Illustrations

All the schemes were created using BioRender.com.

