## supplemental data for "PI3Kδ Bridges Microbial Surveillance with Antigen Presentation to Reinforce Intestinal Immunity"

#### Extended data files

**Extended Data Fig. 1 PI3Kδ inactivation results in the expansion of enteric pathobiont *H. hepaticus*, causing dysbiosis in CONV housing.** (**a**) Lipocalin-2 levels in colon mucus from indicated mouse genotypes, assessed by ELISA (n=5 - 6 per group)., (**b**) Representative H&E-stained section of colon segment from WT-SPF and δD910A-SPF mice, respectively (n=3-4) (scale bars, 100 μM) (**c, d**) RT-PCR analysis from feces of WT-CONV or δD910A-CONV asymptomatic or δD910A-CONV prolapse-presenting mice, (**c**) Total 16S rRNA (n=12), and (**d**)16S *H. hepaticus.* rRNA (n=6-9 per group). (**e**) ELISA quantification of TNF, IL-1β, IL-12p40, IFN-γ, IL-17, and IL-10 levels in medium secreted from unstimulated or ex vivo stimulated (LPS 100 ng/ml for 18 h) colon explants from WT-CONV and δD910A-CONV asymptomatic mice (n=9-10, three independent experiments). Data are presented as means ± SEM. One-way or two-way ANOVA with Tukey's post hoc tests were used; *p* values were considered **p*<0.05, ***p*<0.01, ****p*<0.001, and **** *p*<0.0001.

**Extended Data Fig. 2**. **Systemic PI3Kδ inactivation dysregulates T cell immune balance in mLNs and leads to T cell-mediated colon inflammation.** (**a-b**) $CD4^+$ and $CD8^+$ T cell distribution in mLNs of (**a**) WT or δD910A (**b**) mice, assessed by flow cytometry (n=8 per group). (**c**) The proportions of $CD4^+$ T cells expressing FOXP3, IL-10, or IL-10 IFNγ isolated from cLP of WT, δD910A (*upper* panels) and WT, p110$\delta^{\Delta CD11c}$ (*lower* panels) mice after PMA and ionomycin restimulation *in vitro* (n=5-6). (**c**) Kaplan-Meier analysis of age-dependent development of rectal prolapse in CONV reared *Rag2*$^{-/-}$, δD910A and *Rag2*$^{-/-}$ δD910A mice (n=10-20) ***p<0.0001. Log-rank (Mantel-Cox). (**d**) Representative H&E-stained section of colon segment of *Rag2*$^{-/-}$, δD910A and *Rag2*$^{-/-}$ δD910A mice (n=4-7). (**e**) Comparative DAI scores of mice in *d*. (**g-i**) DSS-induced colitis in WT and δD910A mice in SPF or CONV housing with their Veh-treated controls (n=7-20 mice per group). (**g**) Body weight change (n=12 per group) (**h**) colon lengths (n=6 - 9 per group), and (**i**) comparative DAI (n=4 - 6 per group), (**j**) $Ly6C^+$ myeloid cells in cLP of p110$\delta^{fl/fl}$ and p110$\delta^{\Delta CD11c}$ mice (n=5-6 per group). (**k, l**) p110$\delta^{fl/fl}$ and p110$\delta^{\Delta Lyzm}$ mice were exposed to 2.5% w/v DSS or Veh in drinking water for 5 days. (**k**) Percent change in body weight (N=3, n=3-5) and (**l**) representative images of colons (*left*) and bar charts (*right*) of indicated mouse colon lengths (n=5-9)**.** Data are expressed as means ± SEM. Two-way ANOVA with Tukey's post hoc test or student t-test (unpaired) was performed for statistical analysis, *p* values were considered **p*<0.05, ***p*<0.01, *** *p*<0.001, and *****p*<0.0001.

**Extended Data Fig. 3 PI3Kδ inactivation hyperactivates inflammasome pathway in BM-DCs.** (**a)** Representative flow cytometry histogram plot (*left* panel) and mean fluorescence intensity (MFI) (*right* panel) of MDP-FITC uptake at indicated time intervals in WT or δD910A BM-DCs (n=3). (**b-c**) BM-DCs were rested in medium or activated by LPS (100 ng/ml) or MDP (30 μg/ml) for 18 h. Bart charts showing (**b**) mean fluorescence intensity (MFI) of MHC-II, CD40, and CD86 surface expression analyzed by flow cytometry (n=3), and (**c**) LDH released in the culture supernatants, detected by ELISA (n=5). (**d**) Colony forming units (CFU) counts of live *Salmonella typhimurium* (S.t.), *Staphylococcus aureus* (*S.a.*), and *Pseudomonas aeruginosa* (*P.a.*) in WT or δD910A BM-DCs at 2 h post-infection (n=5 per group). (**e**) Bart charts represent MFI of *E. coli*-FITC phagocytosed at the indicated temperatures in WT and δD910A BM-DCs at 1 h, measured by flow cytometry (n=3). (**f, g**) Representative immunoblot of cleaved caspase-1 (p20 fragment) in WT and δD910A BM-DCs stimulated with (**f**) LPS (100 ng/ml) until 90 min and (**g**) LPS (100 ng/ml) or LPS and ATP (5 mM) for 30 min. All immunoblot images are representative of three independent experiments. (**h**) Mitochondrial (mt)ROS was measured by flow cytometry using mitoSOX$^{Red}$ probe (10 μM) in WT and δD910A BM-DCs stimulated with MDP, LPS, or rested in medium (n=3). Data are expressed as means ± SEM. Two-way ANOVA with Šídák's post-hoc test was performed for statistical analysis; *p* values were considered *$p<0.05$, **$p<0.01$, *** $p<0.001$, and **** $p<0.0001$.

**Extended Data Fig. 4**. **DC-specific PI3Kδ does not impact OVA peptide presentation and T cell proliferation.** (**a, b**) Flow cytometry analysis of OT-II T cells following co-cultures with WT or δD910A BM-DCs preloaded with OVA-coated beads (1:50), containing indicated concentrations of OVA, (**a**) OT-II T cell division, (**b**) The proportions of IL-10$^+$, FOXP3$^{+,}$ and IFNγ$^+$ in divided OT-II T cells after co-cultures with indicated genotypes of BM-DCs as in *a*, followed by PMA and ionomycin restimulation (n=6-9 per group). (**c, d**) OT-I or OT-II T cell division assessed by flow cytometry and (**c**) histograms of OT-I T cells and (**d**) proportions of dividing OT-I or OT-II T cells after co-cultures with WT or δD910A BM-DCs preloaded with indicated concentrations of (**a**) SIINFEKL or (**b**) OVA (323-339) peptides. (n= 2-4). (**c-e**) Mean fluorescence intensity (MFI) of (**c**) 3 μM FITC-bead (1:10) phagocytosis, (**d**) Dextran-FITC (70 kDa, 1 μg/ml) macropinocytosis, and (**e**) BSA (1 μg/ml) endocytosis in WT and δD910A BM-DCs at indicated times and temperatures, analyzed by flow cytometry. Two-way ANOVA with Šídák's or Tukey's post-hoc test was performed for statistical analysis, and *p* values were considered *$p<0.05$, **$p<0.01$, ***$p<0.001$, and **** $p<0.0001$.

**Extended Data Fig. 5**. **PI3Kδ drives NOX2-mediated oxidative burst following PAMP-coupled particle uptake.** (**a**) Intracellular ROS detected with $H_2$DCFDA in WT, δD910A, and *Cybb*$^{-/-}$ BM-DCs pretreated with PI3Kδ inhibitors, PI-3065 (1 μM) or CAL-101 (5 μM) or Veh, followed by zymosan (5 μg/ml) stimulation for 120 min (n=3). (**b**) Mean fluorescence intensity (MFI) of zymosan-FITC (5 μg/ml) phagocytosis in WT and δD910A BM-DCs at 15 min at the indicated temperatures by flow cytometry. (**c**) Colony forming units (CFU) levels representing live *Salmonella Typhimurium* (*S.t.*) and *Staphylococcus aureus* (*S.a.*) in WT and δD910A BM-DCs at 6 h post-infection (n=4-5). (**d**) The mean fluorescence intensity (MFI) of phagosome acidification in WT and δD910A BM-DCs following phagocytosis of *E. coli*-pHrodo$^{red}$ at 37$^{0}$C at 30 and 60 min by flow cytometry (n=3-4). (**e**) Mean fluorescence intensity (MFI) of soluble DQ-OVA proteolysis in WT and δD910A BM-DCs at indicated time intervals assessed by flow cytometry (n=3). Results are expressed as means ± SEM. Two-way ANOVA with Tukey's or Šídák's post hoc test was performed for statistical analysis; *p* values were considered as **p*<0.05, ***p*<0.01, *** *p*<0.001, and ****p*<0.0001.

**Extended Data Fig. 6**. **PRR-coupled tyrosine kinase signaling coordinates PI3Kδ-RAC2 activities.** (**a**) Representative immunoblot (left panel) and quantification (right panel) for p-p38 and p38 MAPK in WT, *Rac2*$^{-/-}$ BM-DCs, or BM-DCs pretreated with EHT1864 (0.5 μM) or Veh (DMSO) for 2 h before stimulation with MDP (30 mg/ml) or LPS (100 ng/ml). (**b**) Representative immunoblot for p-Akt$^{Ser473}$ in BM-DCs pretreated with pan-tyrosine kinase inhibitors, AG-17 (0.5 μM), AG-213 (5 μM), or Genistein (10 μM) for 1 h and then stimulated with LPS for 30 min (n=2). (**c**) Total GTP-loading of RAC1/2/3 was quantified using Rac G-LISA activation assay in TCLs of BM-DCs that were pretreated with pan tyrosine kinase inhibitors, AG-17 (0.5 μM), AG-213 (5 μM), and Genistein (10 μM) for 1 h and then stimulated with LPS for 30 min (n=5). (**d**) Representative immunoblots showing recombinant p110β, but not p110α or p110δ binding to Rac1 GTP[S] in a cell-free system. Total recombinant His-tagged p110 isoforms or His-tagged RAC2 levels were visualized using anti-His mAb. One representative experiment out of two is shown. (**e**) Representative immunoblot showing RAC2 in a complex with p110δ-CAAX but not WT-p110δ following co-immunoprecipitation of p110δ using c-Myc mAb. Immunoblots were visualized using an anti-RAC2 Ab. The immunoblot images are one representative out of two independent experiments. (**f**) Cartoon illustrating PI3Kδ coordination of immune responses in DCs: Linking pathogen detection to antigen presentation. Illustration created using BioRender.com. Data are expressed as means ± SEM. Two-way ANOVA with Šídák's post hoc test was performed for statistical analysis, *p* values were considered as **p*<0.05, ***p*<0.01,****p*<0.001 and *****p*<0.0001.

#### Supplementary data Table S1: Colitis Scoring

| **Disease activity index of mice** | |
|---|---|
| Weight loss | 0: 0 %<br>1: 1-5 %<br>2: 6-15 %<br>3: 16-20 %<br>4: 21-30%<br>5: > 30% |
| Occult blood/gross bleeding | 0: negative<br>1: slightly positive<br>2: positive<br>3: gross bleeding |
| Stool consistency | 0: normal<br>1: loose<br>2: diarrhoea |

| **Histological score of mouse colons** | | |
|---|---|---|
| Inflammation | 0<br>1<br>2<br>3 | none<br>slight<br>moderate<br>severe |
| Range | 0<br>1<br>2<br>3 | none<br>mucosa<br>mucosa + submucosa<br>transmural |
| Regeneration | 1<br>2<br>3<br>4 | almost complete regeneration<br>regeneration with crypt depletion<br>surface epithelium not intact<br>no tissue repair |
| Crypt damage | 0<br>1<br>2<br>3 | none<br>basal 1/3 damaged<br>basal 2/3 damaged<br>entire crypt and epithelium lost |
| Percent involvement | 1<br>2<br>3<br>4 | 1-25 %<br>26-50 %<br>51-75 %<br>76-100 % |

### Supplementary data Table S2: Gating strategy

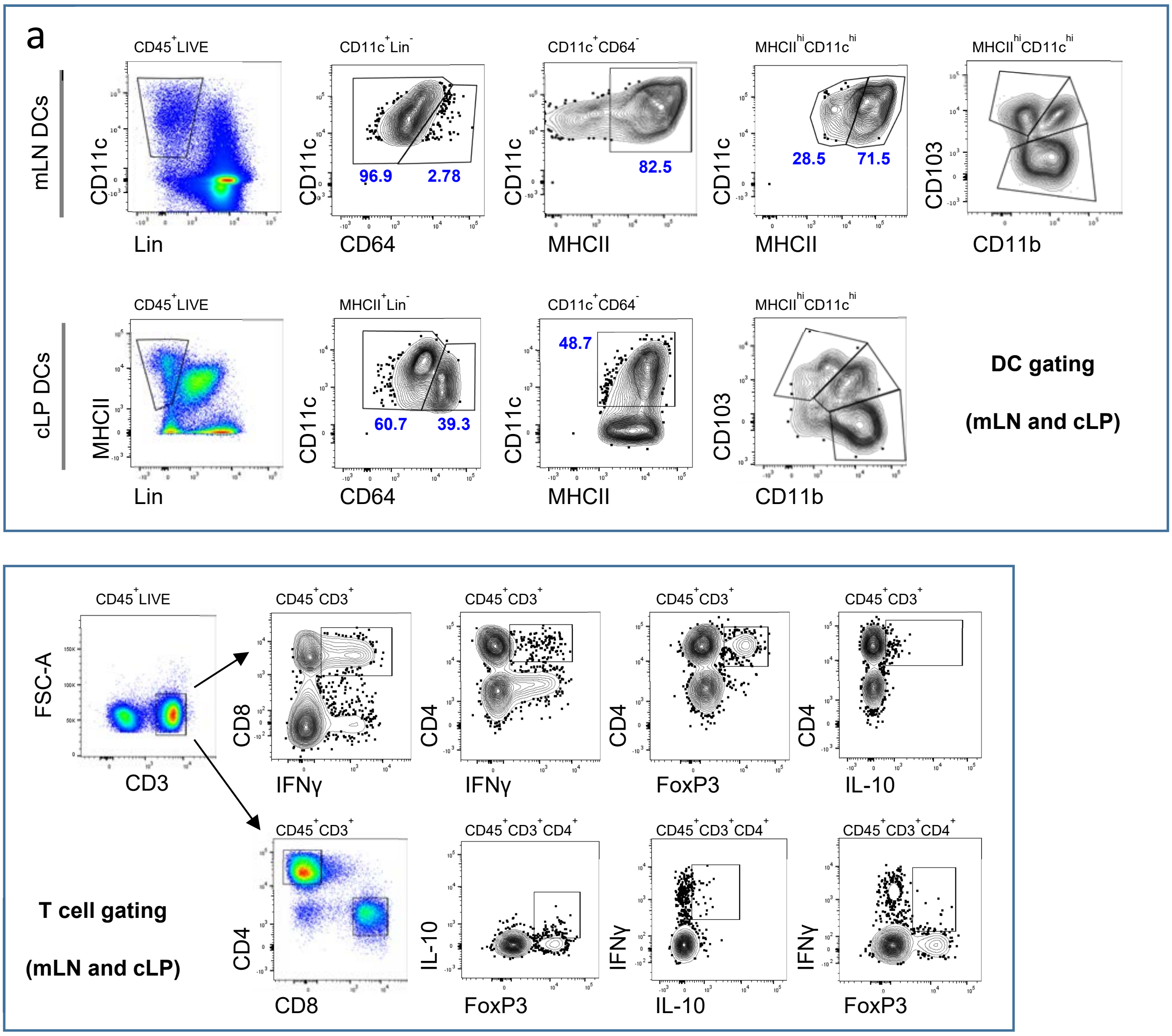


**Supplementary data Table S2. Gating Strategy.** Cells were initially gated based on FSC-A versus SSC-A to identify viable cells. Doublets were excluded using FSC-H against FSC-A. Dead cells were excluded using a live/dead viability marker, and only $CD45^+$ immune cells were included in the analysis across all samples. (**a**) In mLNs, $CD11c^+Lin^-$ cells were gated and then macrophages were excluded based on their CD64 expression. DCs were identified using CD11c and MHC class II markers as $CD11c^{hi}MHCII^{hi}$ and then were further classified using the CD11b and CD103 surface markers. In cLPs, the $Lineage^+$ cells were excluded and DCs were gated as $MHCII^+$. Then, macrophages were excluded based on their CD64 expression. DCs were identified as $MHCII^+$ and $CD11c^+$ ($CD11c^{hi}MHCII^{hi}$) and then were further separated using the CD11b and CD103 surface markers on the cells. (**b**) The $CD3^+$ cell population was gated and separated into $CD4^+$ or $CD8^+$. These cells were considered IFNγ, FOXP3 or IL-10 positive when they fell into the designed gate as shown in the representative image.

| Supplementary data Table S3. KEY RESOURCES | | |
|---|---|---|
| **REAGENT** | **SOURCE** | **IDENTIFIER** |
| **Antibodies for Flow Cytometry** | | |
| FITC anti-mouse MHC Class II (I-A/I-E) | eBioscience | Clone M5/114.15.2; Cat # 11-5321-82 |
| PE anti-mouse Foxp3 | eBioscience | Clone FJK-16s; Cat # 12-5773-82 |
| APC anti-mouse IL10 | eBioscience | Clone JES5-16E3; Cat # 17-7101-82 |
| eFluor 450 anti-mouse CD4 | eBioscience | Clone RM4-5; Cat # 48-0042-82 |
| PerCP anti-mouse CD45 | Biolegend | Clone 30-F11; Cat # 103130 |
| Brilliant Violet 711 anti-mouse CD3 | Biolegend | Clone 17A2; Cat # 100241 |
| Brilliant Violet 785 anti-mouse CD8a | Biolegend | Clone 53-6.7; Cat # 100749 |
| PE anti-mouse Ly-6G | Biolegend | Clone 1A8; Cat # 127608 |
| Brilliant Violet 421 anti-mouse Ly-6C | Biolegend | Clone HK1.4; Cat # 128031 |
| PE/Cyanine5 anti-mouse CD11c | Biolegend | Clone N418; Cat # 117316 |
| PE anti-mouse CD11c | Biolegend | Clone N418; Cat # 117308 |
| FITC anti-mouse CD11c | Biolegend | Clone N418; Cat # 117306 |
| APC anti-mouse CD11c | Biolegend | Clone N418; Cat # 117310 |
| PE/Cyanine7 anti-mouse CD103 | Biolegend | Clone 2E7; Cat # 121426 |
| Brilliant Violet 785 anti-mouse/human CD11b | Biolegend | Clone M1/70; Cat # 101243 |
| APC anti-mouse/human CD11b | Biolegend | Clone M1/70; Cat # 101212 |
| Brilliant Violet 650 anti-mouse F4/80 | Biolegend | Clone BM8; Cat # 123149 |

| | | |
|---|---|---|
| Alexa Fluor 700 anti-mouse CD25 | Biolegend | Clone PC61;<br>Cat # 102024 |
| FITC Anti-mouse [7/4] | Abcam | Clone 7/4;<br>Cat # ab53453 |
| VioGreen anti-mouse CD45 | Miltenyi Biotec | Clone 30F11;<br>Cat # 130-102-412 |
| FITC anti-mouse IL-17A | Miltenyi Biotec | Clone REA660;<br>Cat # 130-111-856 |
| Zombie Aqua Fixable Viability | Biolegend | Cat # 423102 |
| eFluor 780 Fixable Viability | eBioscience | Cat # 65-0865-14 |
| **Antibodies for Immunoblotting** | | |
| PI3 Kinase p110δ (D1Q7R) | Cell Signaling Technology | Cat # 34050S |
| PI3 Kinase p110α (C73F8) | Cell Signaling Technology | Cat # 4249S |
| PI3 Kinase p110β (S-19) | Santa Cruz Biotechnology | Cat # sc-602 |
| PI3 Kinase p110γ | Cell Signaling Technology | Cat # 4252S |
| PI3 Kinase p85 (19H8) | Cell Signaling Technology | Cat # 4257S |
| Phospho-Akt (Thr308) (244F9) | Cell Signaling Technology | Cat # 4056S |
| Phospho-Akt (Ser473) (D9E) | Cell Signaling Technology | Cat # 4060S |
| Phospho-IKKα (Ser176)/IKKβ (Ser177) (C84E11) | Cell Signaling Technology | Cat # 2078S |
| Phospho-p38 MAPK (Thr180/Tyr182) (D3F9) | Cell Signaling Technology | Cat # 4511S |
| Akt (pan) (C67E7) | Cell Signaling Technology | Cat # 4691S |
| p38 MAPK (D13E1) | Cell Signaling Technology | Cat # 8690S |
| IKKα | Cell Signaling Technology | Cat # 2682S |
| IκBα | Cell Signaling Technology | Cat # 9242S |

| | | |
|---|---|---|
| RAC1 | Cytoskeleton Inc. | Cat # ARC03 |
| RAC2 | Invitrogen | Cat #: PA5-88121 |
| Vinculin | Sigma-Aldrich | Cat # V4139 |
| p40-phox | Merck SA - Millipore | Cat # 07-503 |
| NOXA1/p47phox | Merck SA- Millipore | Cat # 07-500 |
| NOXA2/p67phox | Abcam | Cat # ab175293 |
| Cytochrome b245 Light Chain/p22-phox | Abcam | Cat # ab75941 |
| NOX2/gp91phox | Abcam | Cat # ab80508 |
| GAPDH | Abcam | Cat # ab8245 |
| Caspase-1 p10 (M-20) | Santa Cruz Biotechnology | Cat # sc-514 |
| Caspase-1 (14F468) | Santa Cruz Biotechnology | Cat # sc-56036 |
| Caspase-1 (p20) | AdipoGen | Cat # AG-20B-0042-C100 |
| Caspase-1 (p10) | AdipoGen | Cat # AG-20B-0044-C100 |
| Rabbit polyclonal anti-Ovalbumin (OVA) [EPR27117-90] | Abcam | Cat # ab306591 |
| **Antibodies for Immunoprecipitation** | | |
| PI3 Kinase p110α (C73F8) | Cell Signaling Technology | Cat # 4249 |
| PI3 Kinase p110β (S-19) | Santa Cruz + made in-house[35] | Cat # sc-602 + made in-house[35] |
| PI3 Kinase p110δ (D1Q7R) | Cell Signaling Technology | Cat # 34050 |
| FLAG® M2 | Sigma Aldrich | Cat # F3165 |

| | | |
|---|---|---|
| c-Myc (9E10) | Invitrogen | Cat # MA1-980 |
| **PRR Ligands, Chemicals, Probes, Recombinant Proteins, and Cell Culture Reagents** | | |
| Lipopolysaccharides (LPS) from Escherichia coli O55:B5 | Sigma-Aldrich | Cat # L4524 |
| NOD2 Agonist Muramyl dipeptide (L-D isoform, active) (MDP) | InvivoGen | Cat # tlrl-mdp |
| NOD1 Agonist γ-D-glutamyl-meso-diaminopimelic acid (iE-DAP) | InvivoGen | Cat # tlrl-dap |
| Cell wall preparation of S. cerevisiae (Zymosan) | InvivoGen | Cat # tlrl-zyn |
| Heat-killed Salmonella typhimurium (HK-ST) | InvivoGen | Cat # tlrl-hkst2 |
| Adenosine 5'-triphosphate disodium salt (ATP) | InvivoGen | Cat # tlrl-atpl |
| PI3Kδ inhibitor (IC-87114) | Selleckchem | Cat # S1268 |
| PI3Kδ inhibitor (PI-3065) | Selleckchem | Cat # S7623 |
| PI3Kδ inhibitor (Idelalisib, CAL-101, GS-1101) | Selleckchem | Cat # S2226 |
| PI3Kβ inhibitor (TGX-221) | Selleckchem | Cat # S1169 |
| PI3Kα inhibitor (A66) | Selleckchem | Cat # S2636 |
| PI3Kγ inhibitor (AS-252424) | Selleckchem | Cat # S2671 |
| PI3Kα/δ/γ inhibitor (XL147) | Selleckchem | Cat # S1118 |
| Pan class I PI3K inhibitor (GDC 0941) | Axon Medchem | Cat # Pictilisib-1377 |
| EHT-1864 (Rac inhibitor) | Tocris | Cat # 3872 |
| Diphenyleneiodonium chloride (DPI), NADPH oxidase inhibitor | Selleckchem | Cat # S8639 |
| DNase I | Roche | Cat # 10104159001 |
| Collagenase IV | Roche | Cat # 11088858001 |

| | | |
|---|---|---|
| Dispase II | Sigma-Aldrich | Cat # D4693 |
| Phorbol 12-myristate 13-acetate (PMA) | Sigma-Aldrich | Cat # P1585 |
| Ionomycin | Sigma-Aldrich | Cat # I0634 |
| His-tagged Rac2 protein: human wild type | Cytoskeleton Inc. | Cat # RC02 |
| His-tagged Rac1 protein: human wild type | Cytoskeleton Inc. | Cat # RC01 |
| Recombinant human PI3K p110α + PI3K p85 alpha protein | Abcam | Cat # ab196098 |
| Recombinant human PI3K p110β + PI 3Kinase p85 beta protein (Active) | Abcam | Cat # 17180687 |
| Recombinant human PI3K (p110δ/p85 alpha) protein (Active) | Abcam | Cat # ab125633 |
| Gibco™ MEM Non-Essential Amino Acids | ThermoFisher Scientific | Cat # 11140050 |
| Gibco™ L-Glutamine | ThermoFisher Scientific | Cat # A2916801 |
| Gibco™ Penicillin-Streptomycin | ThermoFisher Scientific | Cat # 15140122 |
| Gibco™ Sodium Pyruvate | ThermoFisher Scientific | Cat # 11360070 |
| Fetal Bovine Serum, certified | ThermoFisher Scientific | Cat # 16000044 |
| Protein Transport Inhibitor Cocktail | ThermoFisher | Cat # 00-4980-93 |
| Protease and Phosphatase inhibitors | ThermoFisher | Cat # 78442 |
| RBC Lysis Buffer | eBioscience | Cat # 00-4333-57 |
| CD11c MicroBeads UltraPure | Miltenyi Biotec | Cat # 130-108-338 |
| Recombinant Murine GM-CSF | PreproTech | Cat # 315-03 |
| DQ™ Ovalbumin | Invitrogen | Cat # D12053 |

| | | |
|---|---|---|
| EndoFit Ovalbumin | InvivoGen | Cat # vac-pova |
| OVA 323-339 | InvivoGen | Cat # vac-isq |
| OVA 257-264 | InvivoGen | Cat # vac-sin |
| DCFDA / H2DCFDA | Abcam | Cat # ab113851 |
| Phorbol 12-myristate 13-acetate (PMA) | Sigma-Aldrich | Cat # P8139 |
| Green Fluorescent FAM-FLICA® Caspase-1 (YVAD) Assay Kit | 2B Scientific Ltd. | Cat # 98-100TESTS |
| Albumin from Bovine Serum (BSA), Tetramethylrhodamine conjugate | Invitrogen | Cat # A23016 |
| Dextran, Tetramethylrhodamine | Invitrogen | Cat # D1817 |
| Zymosan A BioParticles™, Alexa Fluor™ 488 conjugate | Invitrogen | Cat # Z23373 |
| pHrodo™ Red Zymosan Bioparticles | Invitrogen | Cat # P35364 |
| pHrodo™ Red-E. coli BioParticles | Invitrogen | Cat # P35361 |
| pHrodo™ Red S. aureus Bioparticles | Invitrogen | Cat # A10010 |
| DQ™ Ovalbumin | Invitrogen | Cat # D12053 |
| FluoSpheres™ Amine-Modified Microspheres, 1.0 µm | Invitrogen | F8765 |
| Aliphatic Amine Latex Beads, 2% w/v, 3 µm | Invitrogen | Cat # A37368 |
| Dynabeads™ M-280 Streptavidin | Invitrogen | Cat # 11205D |
| CD11c MicroBeads UltraPure, mouse | Miltenyi Biotec | Cat # 130-125-835 |
| DSS, Dextran Sodium Sulfate | MP Biomedicals | Cat # 02160110 |
| **ELISAs and other kits** | | |

| | | |
|---|---|---|
| Mouse IL-1 beta/IL-1F2 DuoSet ELISA | R&D Systems | Cat # DY401 |
| Mouse TNF-alpha DuoSet ELISA | R&D Systems | Cat # DY410 |
| Mouse IL-10 DuoSet ELISA | R&D Systems | Cat # DY417 |
| Mouse IL-12/IL-23 p40 Allele-specific DuoSet ELISA | R&D Systems | Cat # DY499 |
| Mouse IL-12 p70 DuoSet ELISA | R&D Systems | Cat # DY419 |
| Mouse IFN-gamma DuoSet ELISA | R&D Systems | Cat # DY485 |
| Mouse IL-17 DuoSet ELISA | R&D Systems | Cat # DY421 |
| Mouse Lipocalin-2/NGAL DuoSet ELISA | Cayman | Cat # DY1857 |
| LDH Cytotoxicity Assay Kit | Cayman Chemical | Cat # CAY601170 |
| PowerViral® Environmental RNA/DNA Isolation Kit | Qiagen Ltd. | Cat # 28000 |
| SuperScript™ VILO™ cDNA Synthesis Kit | ThermoFisher Scientific | Cat # 11754050 |
| SYBR™ Green Universal Master Mix | ThermoFisher Scientific | Cat # 4309155 |
| PAK-PBD Beads | Cytoskeleton | Cat # PAK02 |
| **Plasmids** | | |
| pcDNA3-Flag Rac2 | Addgene | Plasmid #12192 |
| pMX-Myc-p110δ-WT | Made in-house | citation[35] |
| pMX-Myc-p110δ-CAAX | Made in-house | citation[35] |

### Extended Figure 1

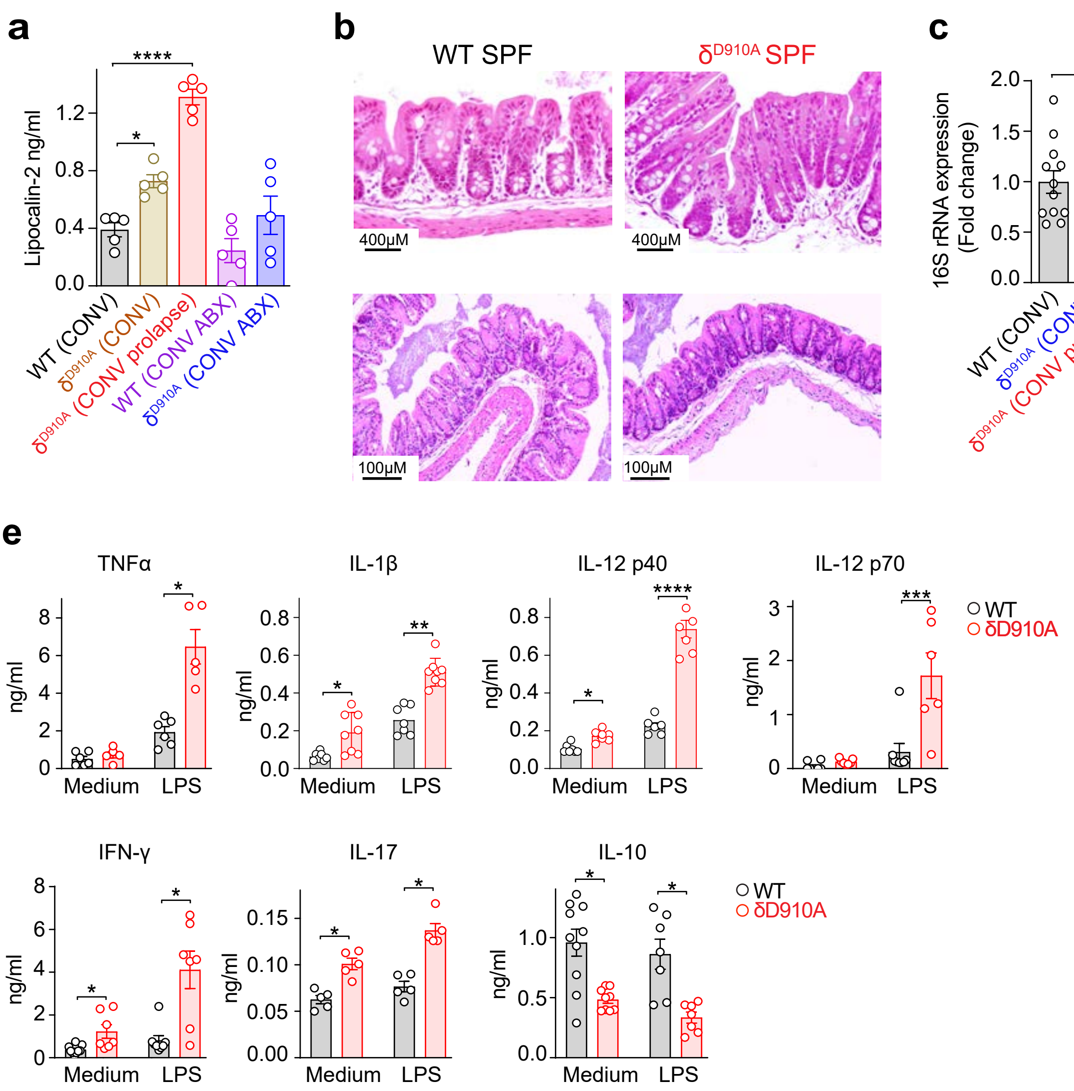

### Extended Figure 2

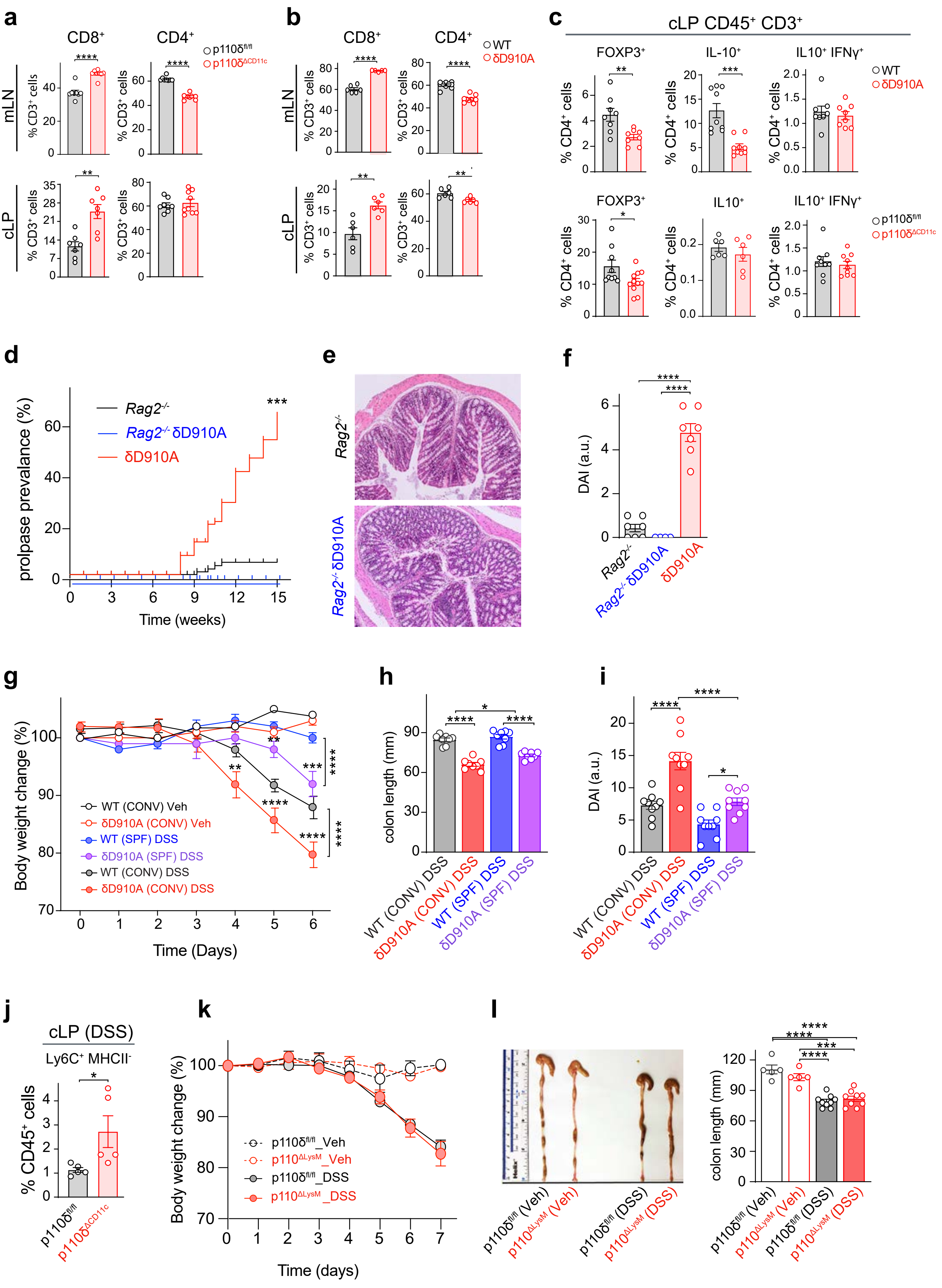

### Extended Figure 3

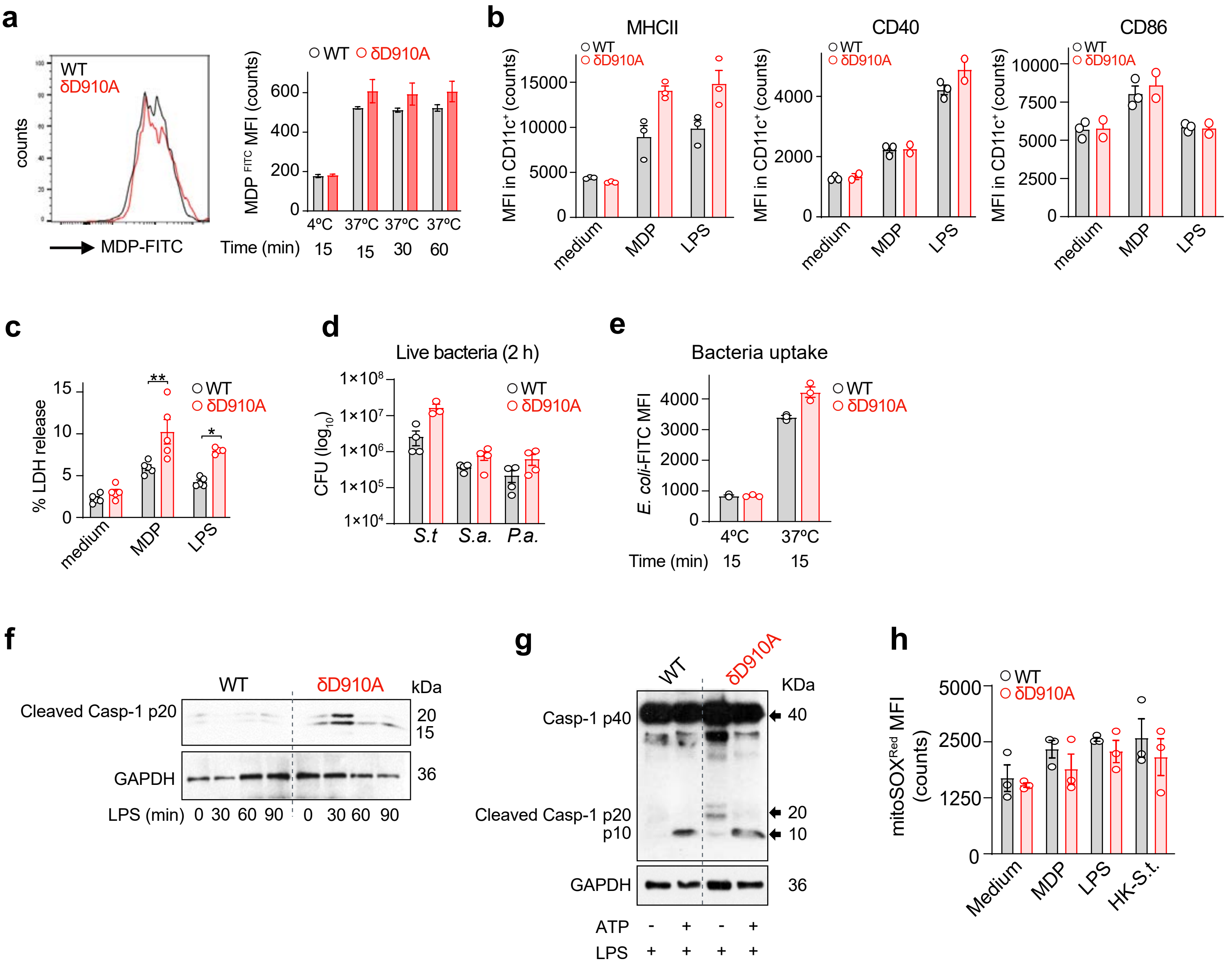

### Extended Figure 4

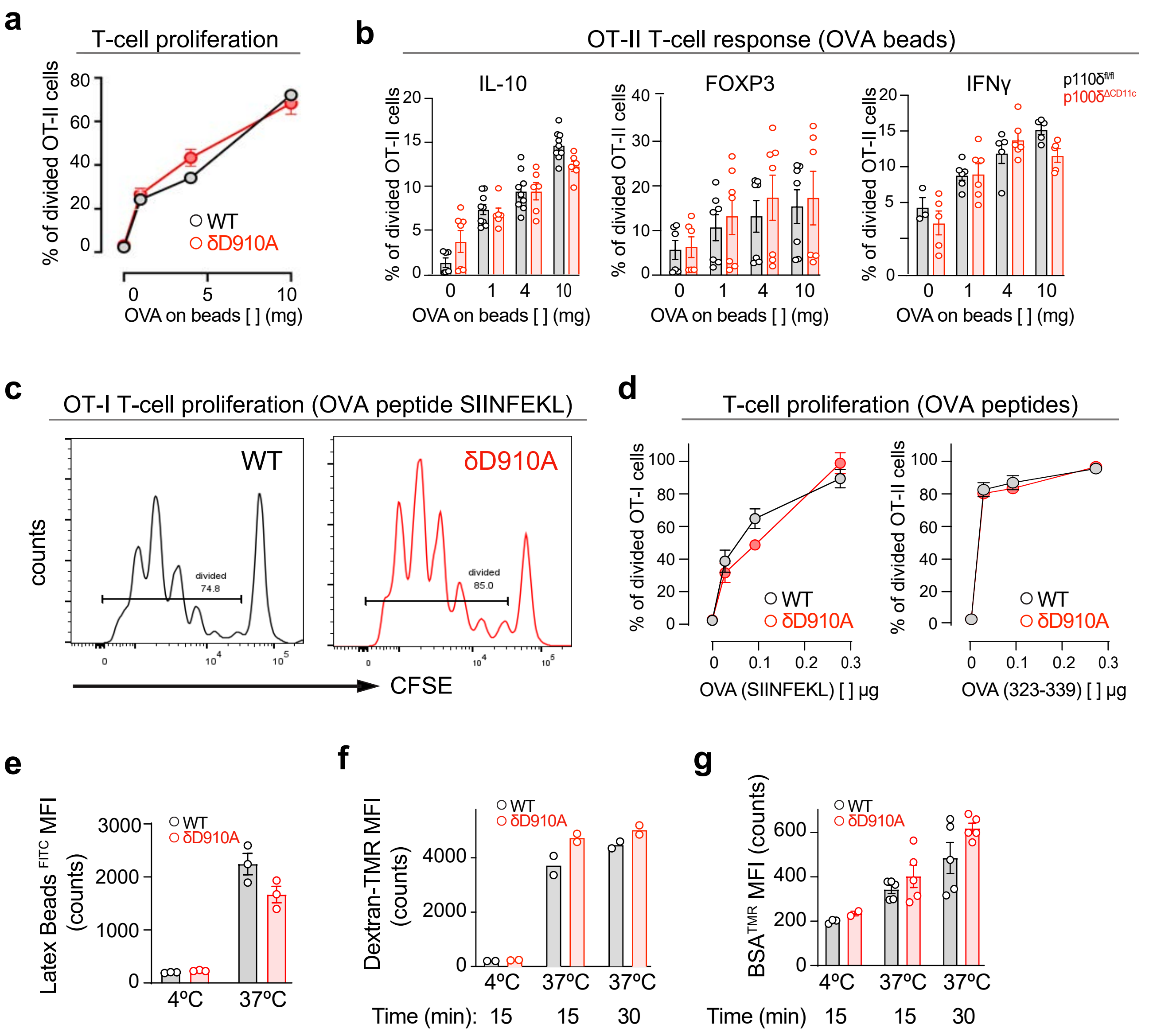

### Extended Figure 5

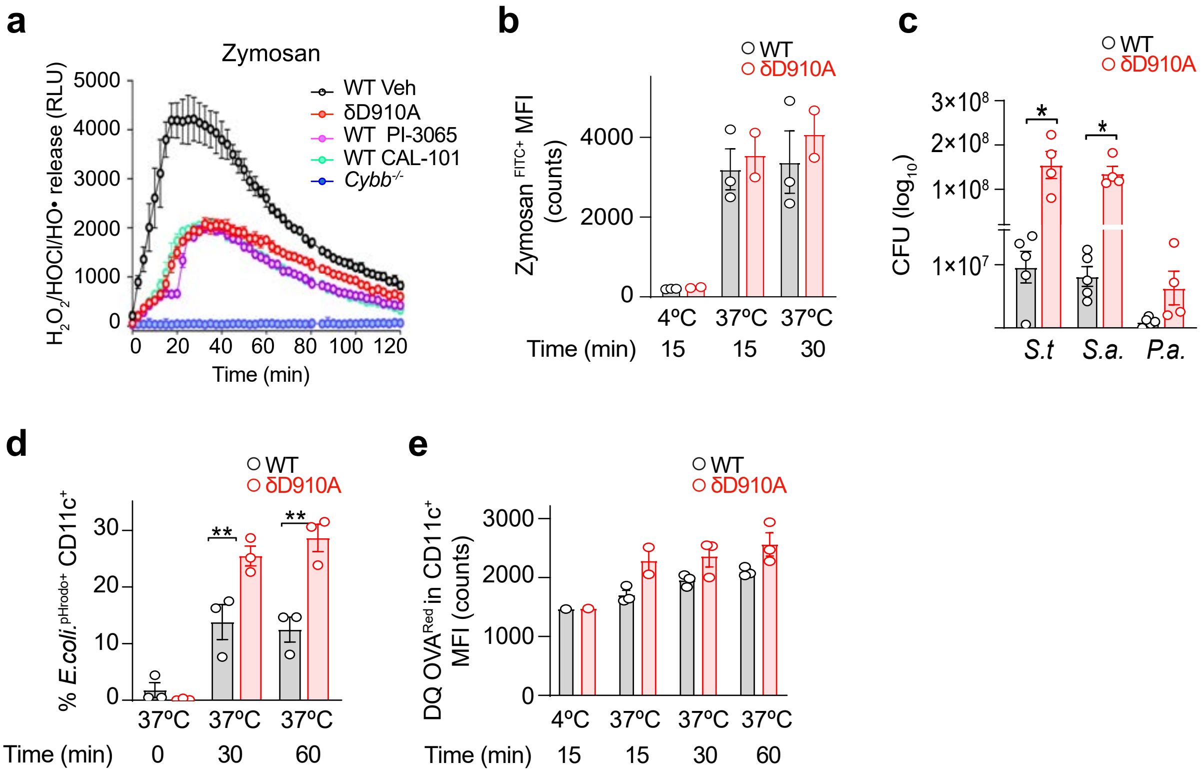

### Extended Figure 6

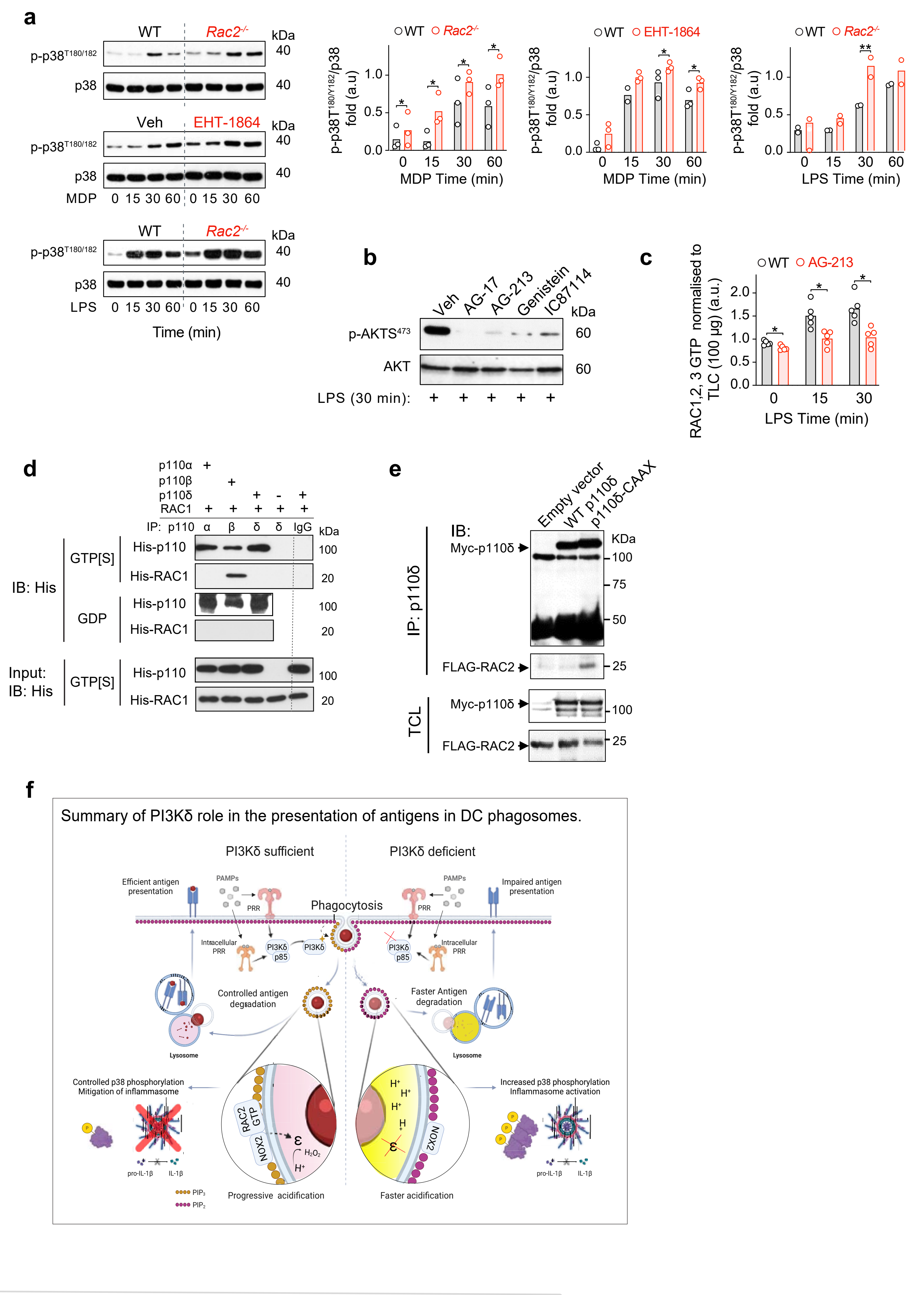
